# Autonomic reflex plasticity associates with time-dependent SUDEP susceptibility in a murine model with hyperreactive stress circuits

**DOI:** 10.1101/2025.11.11.687816

**Authors:** Sandy E. Saunders, Kaylie E. Dow, Grace E. Bostic, Jeffery A. Boychuk, Jamie L. Maguire, Carie R. Boychuk

## Abstract

Sudden unexpected death in Epilepsy (SUDEP) is the leading cause of death in patients with Epilepsy. Although SUDEP results from cardiorespiratory arrest, it’s underlying mechanisms are poorly understood. Considering the significant association between stress-related disorders and Epilepsy, we hypothesized that stress exaggerates autonomic reflexes critical in cardiorespiratory function and that these exaggerated reflexes increase susceptibility to SUDEP. Experiments were performed using a novel mouse model of SUDEP where hyperreactivity of central corticotropin-releasing hormone (CRH) neurons (*Kcc2/Crh*) predisposes mice to SUDEP in the weeks following seizure induction based on the ventral intrahippocampal kainate (vIHKA) model of chronic Epilepsy. In our study, the vIHKA model was employed in both wild-type (WT) and *Kcc2/Crh* mice while they were monitored with EEG and ECG using in vivo telemetry and underwent terminal autonomic reflex testing at time points when mortality peaked and plateaued. A resting tachycardia developed by one week following vIHKA injection but subsided by day 30 in both WT and *Kcc2/Crh* mice. During spontaneous seizures, *Kcc2/Crh* mice had more pronounced reflex-like ictal bradycardias compared to WT controls that notably occurred prior (∼10 sec) to seizure termination. vIHKA injection promoted time-dependent exaggeration of autonomic reflexes, with *Kcc2/Crh* mice exhibiting robust autonomic disturbances compared to WT controls, including a pronounced serotonin-mediated Bezold Jarisch reflex. Taken together, our findings indicate that increased autonomic disturbance burden parallels time-dependent SUDEP susceptibility in mice with hyperreactive stress circuits.

## INTRODUCTION

Sudden unexpected death in Epilepsy (SUDEP) is a significant health burden representing the most common cause of death outside of trauma or accident in patients with Epilepsy (Tomson et al., 2005). Seizure events are often temporally linked to SUDEP whereas more global long-term pathophysiological remodeling is also suspected (Pathak et al., 2025). Cardiorespiratory dysfunction is strongly implicated in SUDEP as severe bradycardia and apnea are observed at seizure termination, or immediately post-ictal, in patients that succumb (Ryvlin, Nashef, Lhatoo, et al., 2013). Similar severe bradycardia and apneas also appear in individuals that eventually succumb to SUDEP (Lamrani et al., 2023), suggesting a predictive causal capacity of these cardiorespiratory disturbances for patients at risk. While these epidemiological factors are being established, diagnostic and interventional patient care for SUDEP remain a critical health need.

Animals models provide a mechanistic approach to understanding how dysfunction of cardiorespiratory control circuits drives clinical presentation and mortality in SUDEP. Most transgenic animal model investigations into mechanisms of SUDEP include mutations known to impact neurodevelopment and intrinsic cardiac excitability (Gu et al., 2024). Stress signaling is an additional important component to Epilepsy and SUDEP. Stress increases the probability of seizure generation and worsens seizure severity (Moore et al., 2014; Si et al., 2024). There is also a significant comorbidity between stress-related disorders and Epilepsy with individuals affected by both succumbing to SUDEP at higher rates than those with Epilepsy alone (Sawyer & Escayg, 2010). These studies implicate the interaction between central stress circuits and autonomic brainstem motor output as a critical mechanism of SUDEP that is distinct from pathological changes in neurodevelopment and intrinsic cardiac excitability.

At the apex of central stress circuits, CRH neurons in the paraventricular nucleus of the hypothalamus (PVN^Crh^) are of particular interest. For example, multiple chemical convulsant models of Epilepsy demonstrate robust, dose dependent, chronic increases in serum corticosterone as a marker of chronic stress activation (Hooper et al., 2018; O’Toole et al., 2014). PVN^Crh^ neuron activity is also significantly elevated by spontaneous seizures across convulsant models as demonstrated by in vivo and in vitro approaches (O’Toole et al., 2014). To link SUDEP with stress and autonomic dysfunction, this study tested mice with hypothalamic–pituitary–adrenal (HPA) axis hyperreactivity via genetic loss of K+/Cl− cotransporter (KCC2) in corticotropin-releasing hormone (CRH) neurons (*Kcc2/Crh* mice). This loss of *Kcc2* in CRH neurons has been previously confirmed and shown to cause an exaggerated HPA axis response to stress that is absent in baseline conditions (Basu et al., 2024; Melon et al., 2018). Chronic Temporal Lobe Epilepsy (TLE) was modeled in *Kcc2/Crh* mice by ventral intrahippocampal kainate administration (vIHKA) (Basu et al., 2024; Zeidler et al., 2018).

In agreement with prior studies, *Kcc2/Crh* mice given TLE by vIHKA showed a dramatic increase in long-term susceptibility to SUDEP thus prompting further mechanistic investigation (Basu et al., 2024). PVN^Crh^ neurons send their densest descending brainstem projection to the cardiorespiratory-regulating nucleus of the solitary tract (nTS) that is the terminal field of most cardiorespiratory sensory afferents (Lee et al., 2013; Moga et al., 1990; Ruyle et al., 2023; Wang et al., 2019). Both isoforms of the CRH receptor are present in nTS (Wang et al., 2018) and excitation of nTS projecting PVN^Crh^ neurons can increase heart rate (HR) (Wang et al., 2019) and modulate cardiorespiratory reflex responsivity (Ruyle et al., 2023). Since abnormal autonomic responsivity is suggested to increase SUDEP susceptibility (Budde et al., 2020; Nakase et al., 2016; Stewart et al., 2017), we hypothesized that hyperreactive CRH neurons in mice with spontaneous seizures results in autonomic reflex disturbances that may underlie SUDEP. To test this, we utilized a combination of chronic electrocardiogram (ECG) and electroencephalogram (EEG) telemetry recordings with acute autonomic reflex testing before and following vIHKA injection in our unique bi-transgenic *Kcc2/Crh* mice compared to WT controls.

## RESULTS

### Resting HR before and after vlHKA injection

To examine the interaction between CRH neuron hyperexcitability and seizures on resting HR, we recorded HR in both *Kcc2*/*Crh* and WT mice using ECG telemetry at various time points following vIHKA injection. Prior to hippocampal injection (Fig. 1A, Table 1.1), there was no difference in resting HR between *Kcc2*/*Crh* and WT mice. However, 7 days after vIHKA injection, HR was elevated in both WT and *Kcc2/Crh* mice (Fig. 1C, Table 1.2). At 14 days post vIHKA injection, HR remained significantly elevated only in *Kcc2/Crh* mice (Fig. 1C, Table 1.2). There were no differences in resting HR among any group on day 21 or 28. To determine if the magnitude of resting tachycardia was predictive of SUDEP susceptibility, we plotted individual vIHKA injected *Kcc2*/*Crh* mice that died, against the average resting HR of vIHKA injected *Kcc2*/*Crh* mice that survived to endpoint (Fig. 1D). We did not observe any qualitative relationship between the magnitude of resting tachycardia and those that died or survived to endpoint, suggesting the severity of resting tachycardia is not predictive of SUDEP susceptibility. Taken together, vIHKA injection induces a resting tachycardia that resolves over time. Further, resting tachycardia does not appear to be predictive of SUDEP but likely contributes to total autonomic disturbance burden.

**Fig 1.**
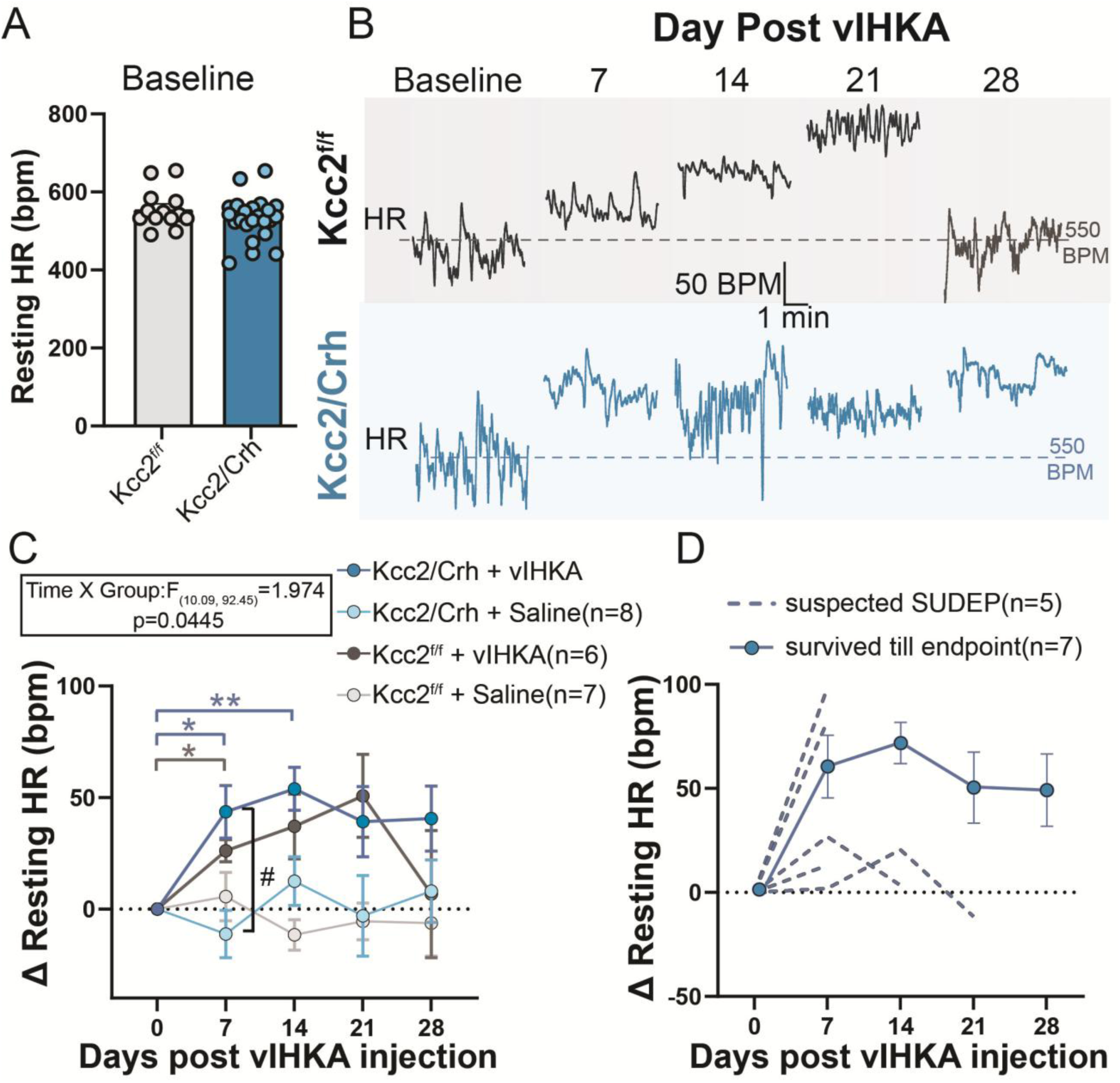
Mice have resting tachycardia shortly after vIHKA injection. A) Resting HR is no different between WT (gray, Kcc2^f/f^) and mice with hyperreactive stress circuits (blue, *Kcc2/Crh*) prior to hippocampal injection. B) Example HR traces from mice with and without hyperreactive stress circuits with telemeters that were monitored weekly following vIHKA injection. C) Resting HR was significantly elevated 7 days after vIHKA injection in both mice with (*p=0.0190, Mixed Effects Analysis and Tukey’s Post Hoc) and without hyperreactive stress circuits (*p=0.0169, Mixed Effects Analysis and Tukey’s Post Hoc). HR remained elevated in mice with hyperreactive stress circuits (**p=0.0017, Mixed Effects Analysis and Tukey’s Post Hoc) but not WT, 14 days after vIHKA injection. D) Mice with hyperreactive stress circuits that died following vIHKA injection are plotted individually (dashed lines). The average HR over time of mice with hyperreactive stress circuits that survive to endpoint are plotted together (solid line). There appears to be no relationship between magnitude of resting tachycardia and SUDEP susceptibility. Symbols are group means ± SEM, *p < 0.05, **p < 0.01, ***p < 0.001 compared to day 0 by mixed effects model and Tukey’s post-hoc test, #p < 0.05 compared to sham control by mixed effects model and Tukey’s post-hoc test.

### Changes in HR during spontaneous seizures

To examine the impact of CRH neuron hyperexcitability on control of HR during seizures, we recorded spontaneous seizures in both *Kcc2*/*Crh* and WT mice using EEG and ECG telemetry recordings (Fig. 2, Table 2.1). In agreement with prior studies (Basu et al., 2024), we observed no difference in average seizure duration between *Kcc2*/*Crh* and WT mice. In both *Kcc2/Crh* and WT mice, HR often (12/14 seizures) increased at ictal start or immediately before seizure onset (Fig. 2D). Average ictal HR was predominately (4/6 WT seizures and 5/8 Kcc2/Crh seizures) increased relative to the preictal period, with no difference between mice with and without hyperreactive CRH neurons.

**Fig 2.**
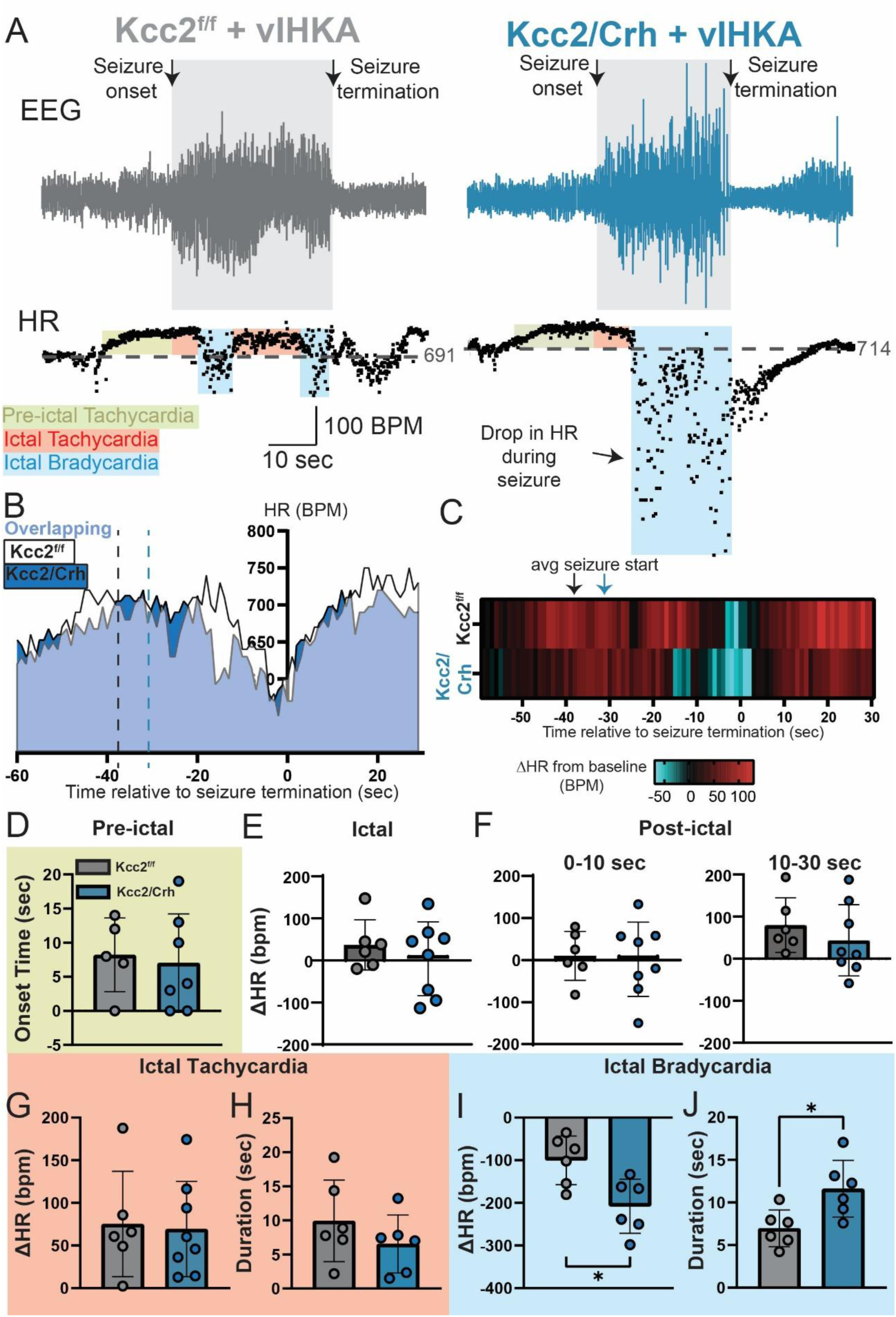
Mice with hyperreactive CRH neurons have greater ictal bradycardia than WT mice. A) Example telemetry recordings of EEG (top) and heart rate (HR, bottom) during a spontaneous seizure in WT (left, gray) and *Kcc2/Crh* (right, blue) mice. Prior to seizure onset, HR began to rise (pre-ictal tachycardia, light green). During the seizure, elevated HR (ictal tachycardia, red) was interrupted by transient drops in HR (blue). *Kcc2/Crh* mice showed severe drops in HR during seizures. B) Qualitative representation of grouped average HR in reference to seizure termination. HR during seizures in WT (white shading) and *Kcc2/Crh* (Blue shading) were similar with *Kcc2/Crh* mice having more pronounced drops in HR right before seizure termination. C) Heatmap depicting grouped average change in HR relative to baseline in reference to seizure termination. *Kcc2/Crh* mice had a more pronounced drop in HR right before seizure termination. D-K, Summary data of mean ictal HR, pre-ictal HR change time of onset, postictal HR 0-10 and 10-30 seconds following seizure termination, ictal tachycardia magnitude, ictal tachycardia duration, ictal bradycardia magnitude, and ictal bradycardia duration. *Kcc2/Crh* mice had significantly greater magnitude (p=0.0264, unpaired t-test) and duration (p=0.0350, unpaired t-test) of ictal bradycardia compared to WT mice. Bars are group means ± SD,* p < 0.05 by unpaired t test.

During seizures (n=11/14), HR fluctuated between periods of stable tachycardia and transient reflex-like bradycardia, with a subset of seizures soley having tachycardia (3/14 seizures). To better quantify nuanced fluctuations in HR during seizures, we segregated ictal HR time series into periods of ictal tachycardia and bradycardia. Magnitude (Fig. 2G) and duration (Fig. 2H) of ictal tachycardia were similar between *Kcc2/Crh* and WT mice, with preictal tachycardia magnitude also exhibiting no difference. However, *Kcc2/Crh* mice had both longer duration (Fig. 2J) and greater magnitude ictal bradycardia (Fig. 2I) than WT. Qualitatively, ictal bradycardia occurred most prominently within seconds prior to seizure termination in both *Kcc2/Crh* and WT mice (Fig. 2B, C). Independent of genotype, HR recovered to baseline levels during the immediate post-ictal period (0-10 sec and 10-30 sec, Fig. 2F).

### Effect of Epilepsy and hyperreactive CRH neurons on baroreflex function

We next sought to investigate potential mechanisms that could underlie increased ictal bradycardia in mice with hyperreactive CRH neurons. As baroreflex is a critical, gold standard reflex for regulating cardiac function (Salah et al., 2025; Schreihofer & Guyenet, 2002), we investigated both high and low pressure baroreflex responsivity at 10 and 30 days after vIHKA injection or in saline-injected sham controls. To investigate reflex changes in HR mediated by high pressure baroreceptors, we infused phenylephrine (PE), an alpha-adrenergic agonist, into the right jugular vein (Fig. 3). PE infusion increased BP equally at all time points across all groups (Fig. 3C, Table 3.1). As expected, PE infusion led to a reflexive decrease in HR (Fig. 3A, B). A modified-Boltzmann equation (Heusser et al., 2021) was used to relate PE-mediated increases in BP to baroreflex-mediated decreases in HR (inset with colored background in example traces Fig. 3A & B). Baroreflex sensitivity i.e., the slope of the resulting curve (Fig. 3E, Table 3.2) was decreased at day 30 post-VIHKA injection relative to day 10, independent of hyperreactive CRH neurons. As such, baroreflex blunting by day 30 decreases baroreflex sensitivity to control levels as a generic, non-genotype specific response to vIHKA injection.

**Fig 3.**
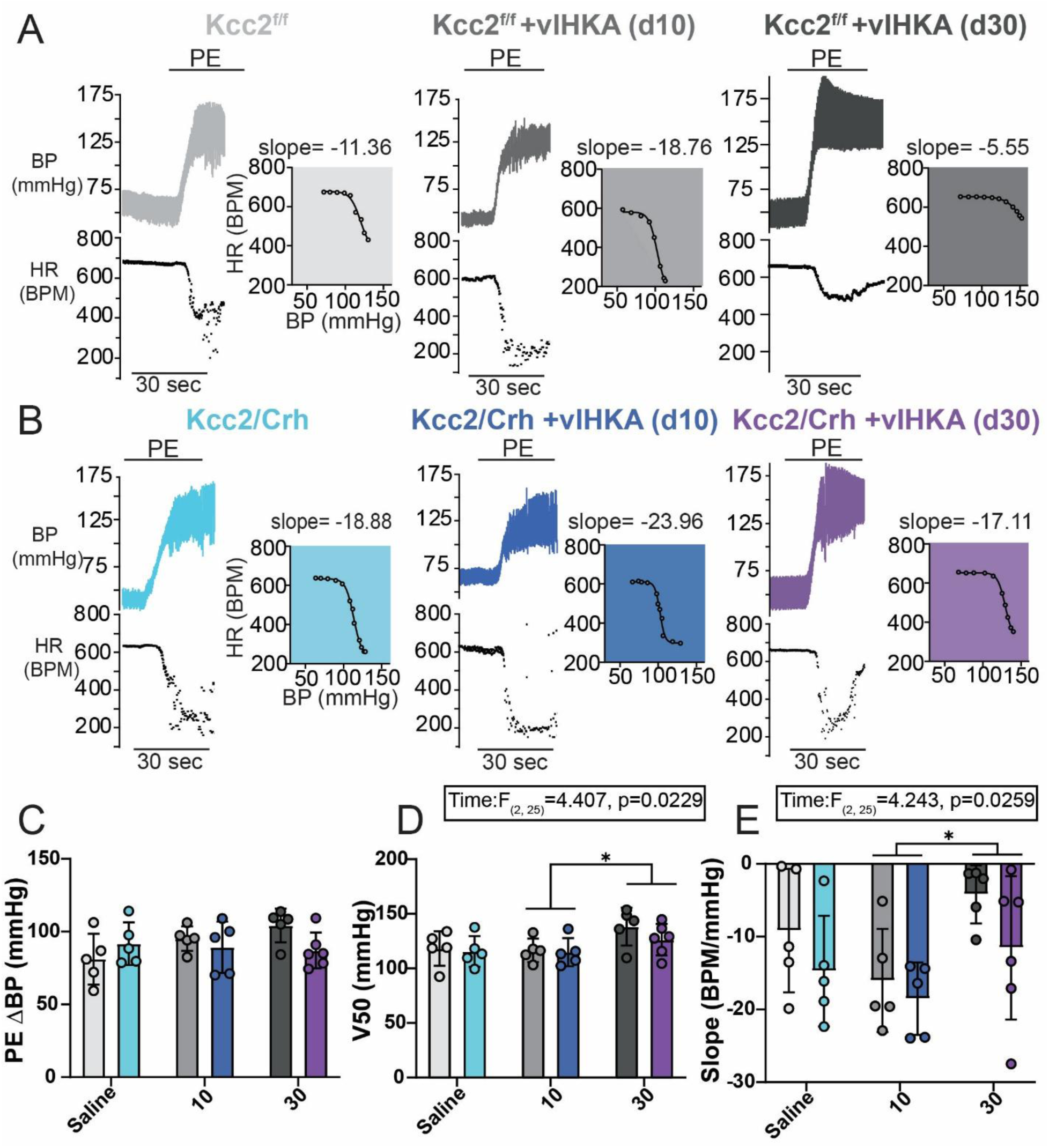
Mice have blunted baroreflex sensitivity 30 days after vIHKA injection. A) Example blood pressure (BP, top) and heart rate (HR, bottom) recordings during jugular infusion of phenylephrine (PE) in WT (*Kcc2^f/f^*) mice at 10 (middle, medium gray) and 30 (right, dark gray) days post vIHKA injection or in saline-injected sham controls (left, light gray). B) Example BP (top) and HR (bottom) recordings during jugular infusion of PE in *Kcc2/Crh* mice either at 10 (middle, dark blue) and 30 (right, purple) days post vIHKA injection or in saline-injected sham controls (left, light blue). In all experiments, infusion of PE caused an increase in BP and drop in HR. C) Summary data of PE-mediated increase in BP. D) Summary data of baroreflex set-point (V50) and E) Summary data of baroreflex sensitivity (slope) that was determined via the slope of resulting curve from fit with modified Boltzmann equation (colored inset). On day 30, baroreflex sensitivity (slope) was significantly blunted (*p=0.0198, Two-way ANOVA with Tukey’s Post Hoc) and V50 (*p=0.0359, Two-way ANOVA with Tukey’s Post Hoc) significantly increased compared to day 10, independent of genotype. Bars are group means ± SD, * p < 0.05 by two-way Anova and Tukey’s post-hoc test.

Similar to baroreflex sensitivity, the operating point (V50), which represents the arterial pressure at which pressure disturbances are effectively buffered by reflex response (Fig. 3D) was increased at day 30 post-vIHKA injection, relative to day 10, independent of genotype. Thus, vIHKA injection nonspecifically increases the operating point of baroreflex at a more chronic phase (30 days) post-induction.

To evaluate low pressure baroreceptors, we infused sodium nitroprusside (SNP), a nitric oxide donor, in the right jugular vein (Fig. 4). SNP infusion caused a consistent and rapid decrease in BP that quickly plateaued in all groups (Table 4.1, Fig. 4C). In contrast to BP, a biphasic response in HR was observed during SNP infusion (Fig. 4D, Table 4.2). During the early phase, HR remained stable in all groups. During the late phase, genotype- and treatment-dependent responses were observed. In saline-injected sham mice, HR qualitatively increased during the late phase in *Kcc2/Crh* (Fig. 4B left) but decreased (e.g. a paradoxical bradycardia) in WT (Fig. 4A left). On day 10 following vIHKA injection, late phase paradoxical bradycardia occurred in both WT (Fig. 4A middle) and *Kcc2/Crh* mice (Fig. 4B middle), with the response greater than respective saline-injected sham controls (Fig. 4D). While all mice continued to exhibit late paradoxical bradycardia at day 30 post vIHKA injection, the magnitude was attenuated relative to day 10 in WT (Fig. 4A right) but not *Kcc2/Crh* animals (Fig. 4B right). Furthermore, paradoxical bradycardia 30 days post-vIHKA injection was greater in *Kcc2/Crh* than WT mice. Taken together, paradoxical bradycardia is exaggerated shortly (10 days) after vIHKA injection in both mice with and without hyperreactive CRH neurons. By day 30, this paradoxical bradycardia is attenuated in WT mice, but remains exaggerated in *Kcc2/Crh* mice. Thus, paradoxical bradycardia is non-specifically exaggerated shortly after vIHKA injection (10 days post) but undergoes attenuated blunting in mice with hyperreactive CRH neurons.

**Fig 4.**
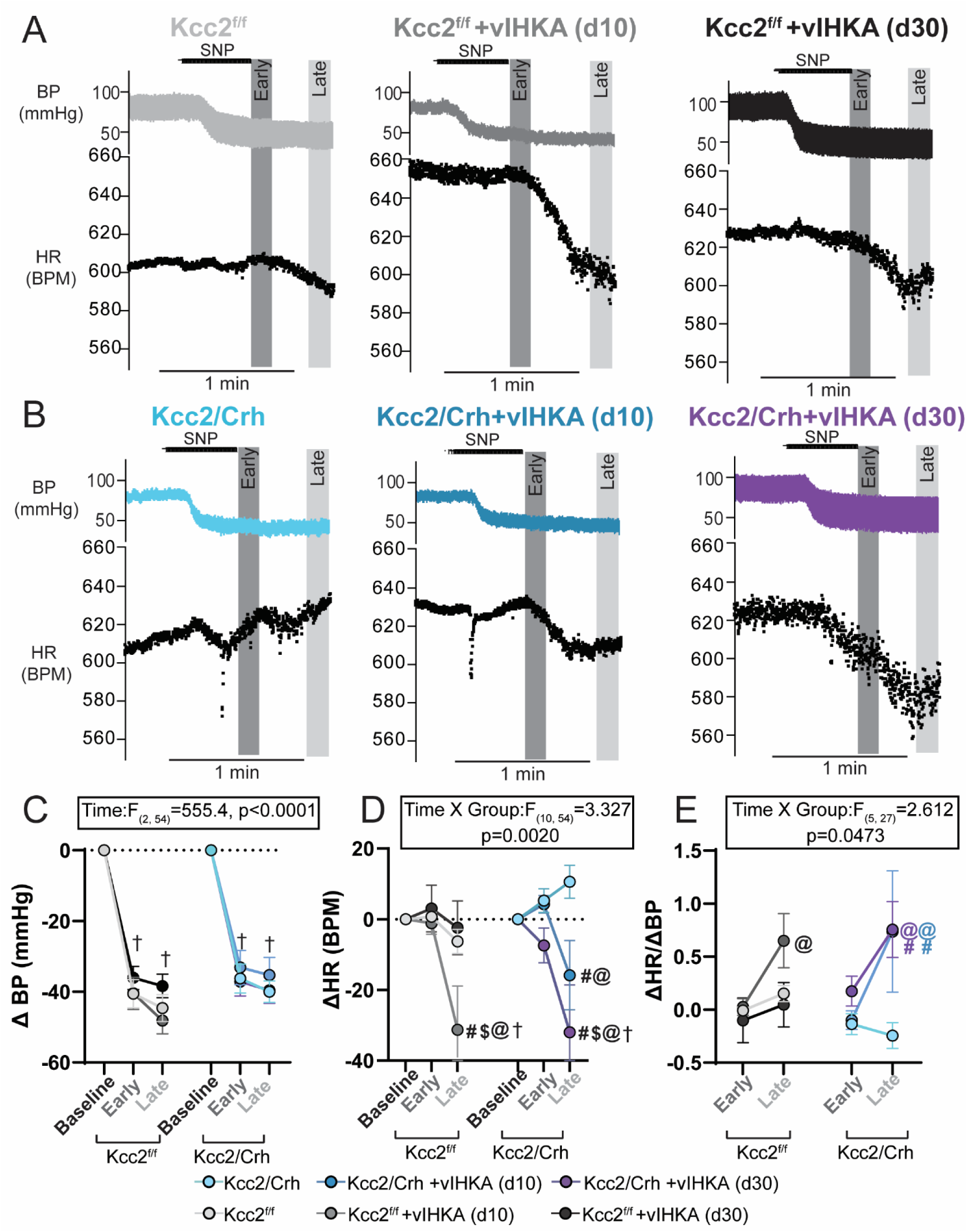
vIHKA injection promotes robust time-dependent changes in paradoxical bradycardia. A) Example blood pressure (BP, top) and heart rate (HR, bottom) recordings during jugular infusion of sodium nitroprusside (SNP) in WT (*Kcc2^f/f^*) mice at 10 (middle, medium gray) and 30 (right, dark gray) days post vIHKA injection or in saline-injected sham controls (left, light gray). B) Example BP (top) and HR (bottom) recordings during jugular infusion of SNP in *Kcc2/Crh* mice either at 10 (middle, dark blue) and 30 (right, purple) days post vIHKA injection or in saline-injected sham controls (left, light blue). In all experiments, infusion of SNP caused a decrease in BP. Changes in HR tended to occur in a biphasic manner, where an initial (early) increase in HR was often followed by a fall in HR (late). C) Summary data showing a significant and sustained drop in BP in collapsed groups following jugular infusion of SNP. D) Summary data of SNP-mediated change in HR, with WT mice having paradoxical bradycardia (significant decreases in HR during late phase) on day 10(@p=0.0004, †p=0.0002, #p=0.0384) and *Kcc2/Crh* mice having paradoxical bradycardia on both day 10(@p=0.0206, #p=0.0346) and day 30(@p=0.0010, †p<0.0001, #p<0.0001). WT mice on day 30 had blunted paradoxical bradycardia relative to WT mice on day 10 ($p=0.0100) and *Kcc2/Crh* mice on day 30 ($p=0.0030). E) Summary data of change in HR over change in BP shows HR changes more for a given change in BP in the late phase on day 10 (@p=0.0106) in WT mice and on day 10 (@p=0.0011, #p=0.0411) and 30 (@p=0.0094, #p=0.0247) in *Kcc2/Crh* mice. Symbols are group means ± SEM, † p < 0.05 compared to group baseline, # p < 0.05 compared to saline-injected sham control in the same phase, @ p < 0.05 compared to “early” in group, $ p < 0.05 compared to late WT d30; Analysis by repeated measures two-way ANOVA and Tukey’s post hoc.

To normalize changes in HR for changes in BP, we plotted change in HR over change in BP in the early and late phase (Fig. 4E, Table 4.3) during the biphasic HR response to SNP infusion. At no time was ΔHR/ΔBP during the “early” phase different between genotype or across treatment. Notably, at day 10 and 30, late phase ΔHR/ΔBP of vIHKA-injected *Kcc2/Crh* mice was significantly greater than ΔHR/ΔBP of saline-injected sham *Kcc2/Crh* controls.

### Effect of Epilepsy and hyperreactive CRH neurons on the BJR

As dysfunction of serotonergic signaling is implicated in the pathophysiology of SUDEP (Richerson & Buchanan, 2011), we investigated the Bezold Jarisch Reflex (BJR) (Fig. 5), a cardioinhibitory reflex that is reliably triggered experimentally by activation of cardiopulmonary vagal afferents containing serotonin type 3 receptors (5HT3R)(Fozard, 1983; Yamano et al., 1995). Accordingly, right jugular administration of phenylbiguanide (PBG), a 5HT3R agonist elicited the expected BJR bradycardia (Fig. 5A & B, Table 5.1) (Verberne & Guyenet, 1992). This PBG-dependent bradycardia was vagally mediated since IP administration of atenolol had no effect on reflex bradycardia, but administration of methylscopolamine eliminated it (Fig. 5A & B, Table 5.1).

**Fig 5.**
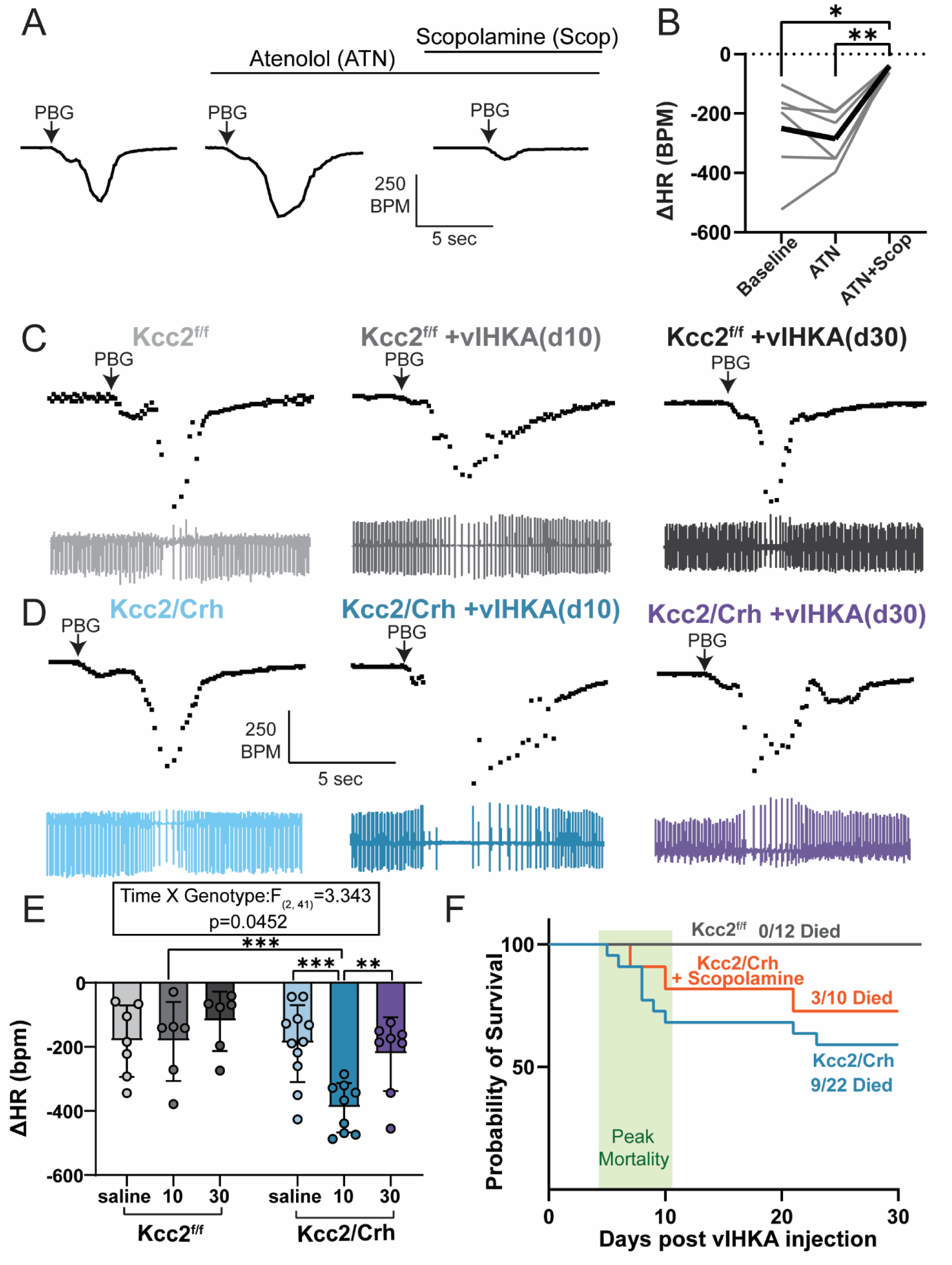
vIHKA injection promotes robust time-dependent increase in PBG-mediated BJR bradycardia in mice with hyperreactive CRH neurons, which contributes to a subset of SUDEP mortality. A) Example trace showing jugular infusion of phenylbiguanide (PBG) elicits the Bezold Jarisch Reflex (BJR), which is characterized by a transient decrease in heart rate (HR). The PBG-mediated BJR bradycardia is not blocked by IP administration of atenolol, but subsequent administration of scopolamine abolishes PBG-mediated BJR bradycardia. B) Summary data (bold line is average; individuals in small gray lines, n=6) showing the PBG-mediated BJR bradycardia is significantly decreased with administration of scopolamine and atenolol compared to atenolol alone (**p=0.0023, Repeated Measures One-way ANOVA with Tukey’s Post Hoc) or baseline (*p=0.0441, Repeated Measures One-way ANOVA with Tukey’s Post Hoc). C) Example ECG traces (bottom) with derived HR (top) during jugular infusion of PBG to elicit the BJR in WT (*Kcc2^f/f^*, gray) at 10 (medium gray) and 30 (dark gray) days post vIHKA injection or in saline-injected sham controls (light gray). D) Example ECG traces(bottom) with derived HR (top) during jugular infusion of PBG to elicit the BJR in mice with hyperreactive CRH neurons (*Kcc2/Crh*) at 10 (dark blue) and 30 (purple) days post vIHKA injection or in saline-injected sham controls (light blue). E) Summary data of PBG-mediated change in HR over time. *Kcc2/Crh* mice have a selective exaggeration of the PBG-mediated BJR bradycardia on day 10 relative to Kcc2/Crh saline-injected sham control (***p=0.0005, Two-way ANOVA with Tukey’s Post Hoc) and WT at day 10 (***p=0.0007, Two-way ANOVA with Tukey’s Post Hoc). This exaggeration is attenuated on day 30 (**p=0.0074, Two-way ANOVA with Tukey’s Post Hoc). F) Following vIHKA injection, 9/22 mice with hyperreactive stress circuits died suddenly, with a peak in mortality occurring about a week post injection. WT mice are not susceptible to SUDEP (0/12 died). Chronic treatment with methylscopolamine attenuated SUDEP mortality (3/10 died). Bars are group means ± SD, *p < 0.05, **p < 0.01, ***p < 0.001.

Saline-injected sham WT and *Kcc2/Crh* mice had similar PBG-mediated BJR bradycardia (Fig. 5C, D left). In WT mice, vIHKA injection had no impact on PBG-mediated BJR bradycardia. However, in *Kcc2/Crh* mice, the magnitude of PBG-mediated BJR bradycardia was significantly increased 10 days post vIHKA injection(Fig. 5D middle) compared to vIHKA-injected WT at day 10 (Fig. 5C middle) and saline-injected sham *Kcc2/Crh* mice(Fig. 5D left). By day 30, however, the increased PBG-mediated BJR bradycardia magnitude in vIHKA-injected *Kcc2/Crh* mice was attenuated (Fig. 5D right). Thus, although baseline BJR function is similar, vIHKA injection led to a selective and transient increase in PBG-mediated BJR bradycardia magnitude in mice with hyperreactive CRH neurons, which is not observed in WT mice.

### Chronic muscarinic blockade reduces SUDEP susceptibility in epileptic mice with CRH neuron hyperactivity

To confirm SUDEP susceptibility in our model, a subset of vIHKA-injected WT and *Kcc2/Crh* mice were monitored for mortality for 28 days. Aligning with prior studies (Basu et al., 2024), 9/22 (41%) vIHKA-injected *Kcc2*/*Crh* mice died with peak mortality occurring around the first week following injection (Fig. 5F blue). In contrast, vIHKA-injected WT mice had no mortality (0/12 mice died) (Fig. 5F gray). To determine the impact of vagal bradycardia on SUDEP mortality, mice were chronically treated with peripherally restricted methylscopolamine. Indeed, muscarinic blockade attenuated SUDEP mortality (3/10, 30%, Fig. 5F orange).

### Changes in hippocampal morphology following vlHKA injection

Hippocampal mossy fiber sprouting (MFS) and dentate granule (DG) cell dispersion are hallmark pathological changes observed in both preclinical models of Epilepsy and in patients with Epilepsy (Sutula & Dudek, 2007). To confirm the severity of these pathological changes were unaffected in *Kcc2*/*Crh* mice (Basu et al., 2024), we performed immunohistochemistry (Fig. 6) to quantify DGCD (Fig. 6B, Table 6.1) and MFS (Fig. 6C, Table 6.2) 10 and 30 days after vIHKA injection or in saline-injected sham controls. Independent of genotype, DG cell dispersion 10 and 30 days post vIHKA injection was increased compared to saline-injected sham controls (Fig. 6B). Additionally, DG cell dispersion was increased at day 30 post vIHKA injection, compared to day 10. In contrast, MFS was increased on day 10 and 30 following vlHKA injection compared to saline-injected sham controls (Fig. 6C), but there was no change in MFS between day 10 and day 30 post vIHKA injection. Taken together, vIHKA injection induces hallmark pathological features of Epilepsy shortly after seizure induction, with DG cell dispersion worsening as Epilepsy progresses in the following weeks, and no effect of genotype detected.

**Fig 6.**
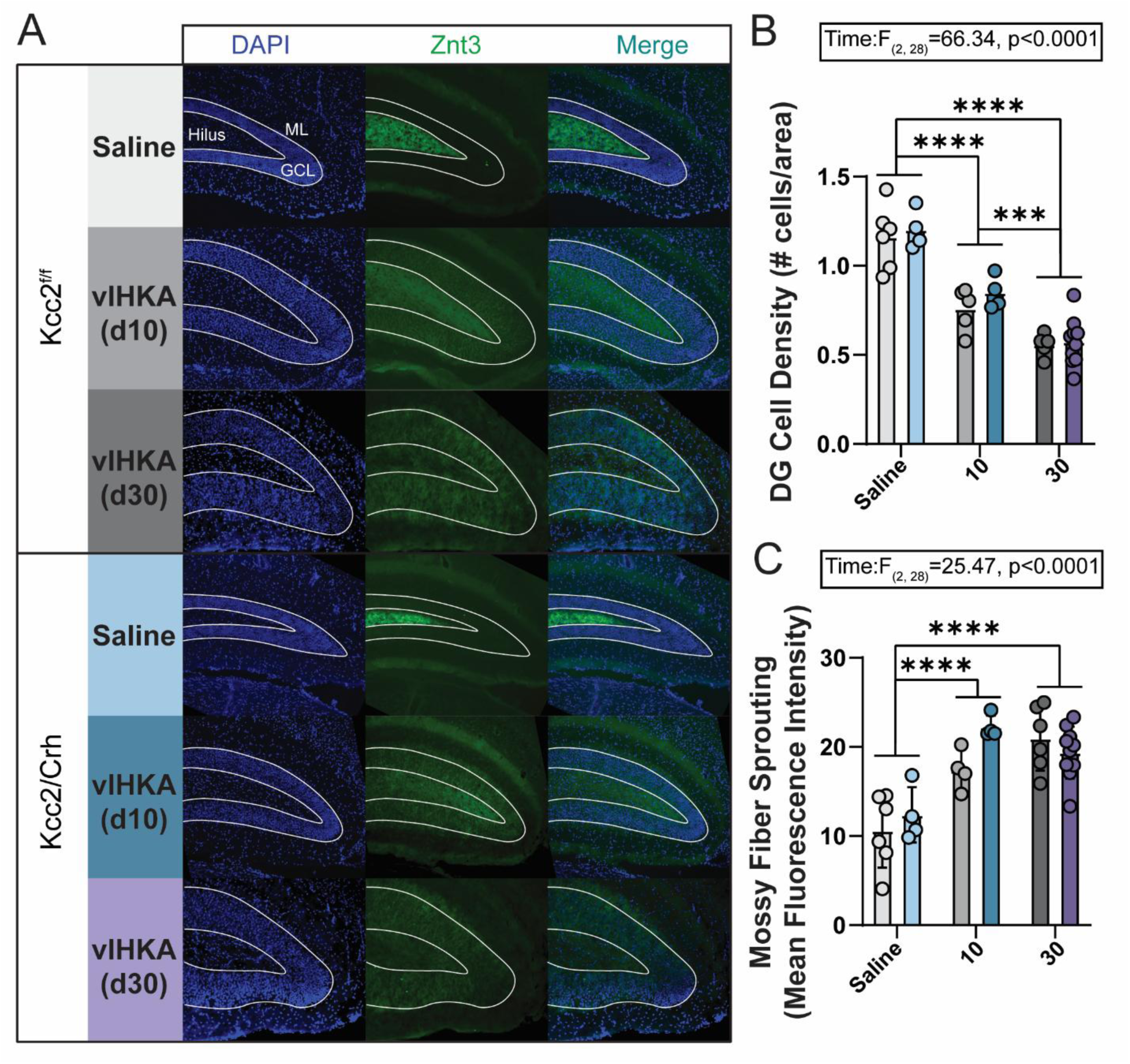
vIHKA injection induces hallmark pathological features of Epilepsy in mice. A) Representative coronal sections of the hippocampus collected from WT mice (Kcc2^f/f^) and mice with hyperreactive stress circuits (Kcc2/Crh) at 10 and 30 days post vIHKA injection, or in sham controls that were injected with saline in the hippocampus. DAPI (blue) stained cell nuclei and was used to quantify dentate granule (DG) cell dispersion (inverse of cell density). Znt3 (green) stained zinc rich mossy fibers which, in normal conditions, project through the hilus. Znt3 was used to quantify mossy fiber sprouting (MFS), or the sprouting of mossy fibers onto other dentate granule cells in the granule cell layer (GCL) and into the molecular layer (ML). B) Summary Data showing a significant increase in DG cell dispersion on day 10 compared to saline-injected sham controls (****p<0.0001, Two-way ANOVA with Tukey’s Post Hoc), independent of genotype. On day 30, DG cell dispersion remained greater than saline-injected sham controls in collapsed groups (****p<0.0001, Two-way ANOVA with Tukey’s Post Hoc). Finally, collapsed groups had significantly increased DG cell dispersion on day 30 compared to day 10 (***p=0.0005, Two-way ANOVA with Tukey’s Post Hoc). C) Summary data showing a significant increase in MFS following vIHKA injection. MFS on day 10 (****p<0.0001, Two-way ANOVA with Tukey’s Post Hoc) and 30 (****p<0.0001, Two-way ANOVA with Tukey’s Post Hoc) was significantly increased compared to saline-injected sham controls, independent of group. Bars are group means ± SD, *p < 0.05, **p < 0.01, ***p < 0.001, ****p < 0.0001 by two-way ANOVA with Tukey’s post hoc.

## DISCUSSION

The present study examined the impact of CRH neuron hyperexcitability on autonomic control of HR in a novel mouse model of SUDEP using chronic EEG and ECG telemetry recordings, terminal autonomic reflex testing, and immunohistochemistry. We found that in the weeks following vIHKA injection both *Kcc2/Crh* and WT mice developed a resting tachycardia that later subsided in both groups (Fig. 1). Regardless of genotype, spontaneous seizures were punctuated by rapid changes in HR, including tachycardic and bradycardic-periods, with robust bradycardias appearing at seizure termination (Fig. 2). Despite no changes in the magnitude of tachycardiac periods, bradycardias during spontaneous seizures in *Kcc2/Crh* mice were more severe compared to WT mice. Finally, vIHKA injection led to time-dependent plasticity of autonomic reflexes, specifically increased PBG-mediated BJR bradycardia and prolonged augmentation of “low pressure” baroreflex in *Kcc2/Crh* mice compared to WT (Fig. 3–5). This exaggerated PBG-mediated BJR bradycardia, and diminished capacity to attenuate paradoxical bradycardia, fit in magnitude and timing to drive increased severity of ictal bradycardia and SUDEP susceptibility. These findings support further investigation into the mechanistic underpinnings of SUDEP including its relationships with stress signaling.

### HR changes over time

In the present study, resting HR was not different between WT and *Kcc2/Crh* mice prior to vlHKA injection (Fig. 1A). However, central administration of CRH (Fisher et al., 1983; Nijsen et al., 2000) and optogenetic stimulation of CRH neurons (Wang et al., 2019) increases HR. Therefore, the present lack of change in resting HR might be inconsistent with expected HR from CRH neuron hyperactivity in *Kcc2/Crh* mice based on these previous acute stimulation studies. A number of factors may be contributing to these HR responses. Most notable, our model utilizes a developmental strategy to knockout *Kcc2* in CRH neurons, which has been confirmed previously (Melon et al., 2018). It remains possible that CRH neurons in *Kcc2/Crh* mice have a compensatory change that minimizes the loss of this gene product. For example, recent evidence implicates excitatory amino acid transporter 3 (EAAT3) function in the establishment of glutamate-dependent GABA synaptic strengthening in PVN neurons (the CRH population most prominently implicated in both autonomic and stress response regulation), which protects PVN from excitotoxicity and is responsible for PVN adaptability (Yamaguchi et al., 2024). It remains possible that a developmental modification in EAAT3 expression could alleviate pressures associated with Kcc2 loss and allow for relative homeostatic regulation of PVN^CRH^ neurons activity. Such homeostatic regulation of PVN^CRH^ neurons would increase their probability of remaining in a “hard off” state with little to no basal firing (Gao et al., 2017; Lovick & Coote, 1988; Savic et al., 2022). This aligns well with lack of elevated plasma corticosterone at baseline in *Kcc2/Crh* mice, compared to WT, because elevated PVN^CRH^ neuron activity should otherwise increase this signal (Basu et al., 2024).

TLE induction in *Kcc2/Crh* mice does induce elevated corticosterone relative to WT mice given this same induction protocol (Basu et al., 2024). Here following vIHKA injection, both *Kcc2/Crh* and WT mice developed a resting tachycardia that resolved in the following weeks. This is consistent with previous work. Homeostatic challenges are known to alter activity of PVN, including activity related to stress axis activation (Palkovits, 2000). In particular, previous work demonstrates that regardless of model or genotype, seizures activate the HPA axis, including increased activation of PVN^CRH^ neurons and elevated corticosterone levels (O’Toole et al., 2014). Therefore, it remains possible that seizure related disturbances are sufficient to overcome any inhibitory mechanisms limiting activity of PVN^CRH^ neurons. In line with no differences in resting HR before vIHKA injection, remodeling may occur in PVN^CRH^ neurons of *Kcc2/Crh* and WT mice following vIHKA injection to return resting HR to baseline by day 30 since number of daily seizures, average seizure duration, or seizure burden does not change during this time (Basu et al., 2024). Although faster interictal HR is observed in patients with various types of epilepsy (Sevcencu & Struijk, 2010), whether this results from epilepsy itself or other factors including treatment with antiepileptic drugs is poorly understood. Accordingly, patients with Epilepsy chronically treated with anti-seizure medication have higher interictal HR than newly diagnosed, non-treated patients, suggesting Epilepsy treatment, and not chronic Epilepsy induces resting tachycardia (Manka-Gaca et al., 2016). Therefore, it remains possible that the resolution of acute tachycardia in patients with Epilepsy is confounded by factors such as antiepileptic drugs.

### HR responses during spontaneous seizures

Ictal tachycardia is a common change in HR reported during seizures (Eggleston et al., 2014). In our study, we observed an increase in HR in both *Kcc2/Crh* and WT mice during spontaneous seizures. This increase in HR began immediately pre-ictal, or directly at seizure onset, aligning with several reports in humans (Mayer et al., 2004; Zijlmans et al., 2002). There was no difference in this seizure-induced tachycardia across genotypes, suggesting it does not play a direct role in the generation of SUDEP mortality. Interestingly, we posit these pre-ictal tachycardias may activate sensory interoceptive afferents (Hsueh et al., 2023) related to increased HR and could play a role in the capacity of some patients with Epilepsy to predict imminent seizures (Mackay et al., 2017).

In both *Kcc2/Crh* and WT mice, this ictal tachycardia during spontaneous seizures was interrupted by transient drops in HR that began prior (∼10 sec) to seizure termination. These ictal bradycardic-episodes were longer with greater magnitudes in *Kcc2/Crh* mice than WT, which aligns with previous work using genetic models of SUDEP (Dhaibar et al., 2019; Paulhus & Glasscock, 2025; Trosclair et al., 2020). In patients with Epilepsy, ictal bradycardia or “ictal bradycardia syndrome” is less commonly reported. As most observed seizures with tachycardia are not generalized tonic clonic seizures (San Antonio-Arce et al., 2024) and generalized tonic clonic seizures are largely associated with SUDEP (Ryvlin, Nashef, & Tomson, 2013), recordings of small subsets of seizures might not be enough to confirm (or reject) ictal bradycardia (Hampel et al., 2017). Critically, robust bradycardic events were confirmed as part of the SUDEP terminal sequence in the MORTEMUS study (Ryvlin, Nashef, Lhatoo, et al., 2013), and primarily associated with temporal lobe seizures (Britton et al., 2006) and link to increased SUDEP risk (Rugg-Gunn et al., 2004). Patients with epilepsy and observed ictal bradycardia are also at a 40% risk of presenting with ictal bradycardia in subsequent seizures, suggesting that robust bradycardias are a consistent pathology in a subset of patients (Hampel et al., 2017). Taken together, it is conceivable that increased severity of ictal bradycardia in *Kcc2/Crh* mice promotes SUDEP susceptibility and could serve as a biomarker for SUDEP risk in patients with Epilepsy. Targeted intervention to blunt ictal bradycardia may hold promise in reducing SUDEP.

### Baroreflex sensory afferent regulatory circuits as potential mechanism for poor HR control in Kcc2/Crh mice

Although high pressure baroreflex blunting can be found in patients with temporal lobe Epilepsy (Dutsch et al., 2006), there is no consistent predictable impact of Epilepsy on BP (Nass et al., 2019). Similarly, our present study found only a mild blunting of “high pressure” baroreflex (e.g. reduced sensitivity and higher set point) 30 days after vIHKA injection. This is consistent with other animal models of seizures where limited or no change in baroreflex is observed, including an audiogenic seizure model (Fazan et al., 2011) and amygdala kindling (Kaya et al., 2005). While differences between human patients and animal models could be species dependent, the chronic use of anti-seizure mediations in patients with Epilepsy confounds modulation of BP (Persson et al., 2003). Whether reduced baroreflex sensitivity in patients with Epilepsy occurs from Epilepsy itself or results from anti-seizure medication should be further explored. Regardless, minor “high pressure” baroreflex blunting here was not specific to *Kcc2/Crh* mice and therefore is unlikely to be a major contributing factor for the development of SUDEP.

While infusion of phenylephrine elicits responses related to “high pressure” baroreceptors, infusion of sodium nitroprusside results in hypotension and blood pooling that when sufficiently severe stimulates “low pressure” baroreceptors located in the atria, ventricles, and pulmonary vessels. Activation of these interoceptive afferents produces a “paradoxical” bradycardia (Heesch, 1999; Kuipers et al., 1984; Oberg & Thoren, 1972) that is not blocked by serotonin receptor antagonist and thus is distinct from BJR (Little et al., 1995). This paradoxical phenomenon mirrors HR responses during the decompensated phase of hemorrhagic shock (Little et al., 1995; Schadt, 1989; Souza et al., 2022) and is common in hypotensive trauma patients (Demetriades et al., 1998). In our experiments, vIHKA injection produced severe paradoxical bradycardia in both WT and *Kcc2/Crh* mice 10 days post injection. The magnitude of this response is corrected only in WT mice by day 30; and therefore, this robust paradoxical bradycardia could be a contributing factor for the cardiac arrest seen during SUDEP. Since WT mice present with paradoxical bradycardia 10 days post vIHKA injection, but do not exhibit SUDEP events, the system’s ability to resolve increased “low pressure” baroreflex circuit sensitivity may be an important step in cardiorespiratory collapse with SUDEP.

### Are peripheral serotonin-dependent sensory afferent circuits (i.e. BJR) a mechanism for robust ictal bradycardias in Kcc2/Crh mice?

Although serotonergic signaling is theorized to play a significant role in SUDEP (Richerson & Buchanan, 2011), this study is one of the first, to our knowledge, to examine the role for peripheral serotonergic signaling in SUDEP-related cardiorespiratory dysfunction. First reported in 1867, activation of cardiopulmonary sensory vagal afferents containing serotonin type 3 receptors (5-HT3Rs) produce a triad of physiological responses consisting of bradycardia, apnea, and hypotension known as the Bezold Jarisch Reflex (BJR) (Cramer, 1915; Jarisch & Richter, 1939). Not only are these the triad of cardiorespiratory events seen during SUDEP (Bozorgi et al., 2013; Ryvlin, Nashef, Lhatoo, et al., 2013), an exaggerated BJR can trigger cardiorespiratory collapse (Jackson et al., 2024; Ou et al., 2004; Poulsen & Jellinge, The Bezold-Jarisch reflex and anaesthesia./2024; So et al., 2013). Critically, *Kcc2/Crh* mice, but not WT, had time dependent exaggeration of the PBG-mediated BJR bradycardia that peaked when population mortality was highest and subsequently attenuated when population mortality plateaued. Although our report is the first to link BJR to SUDEP, serum serotonin levels are elevated following generalized seizures (Murugesan et al., 2018), likely via release from activated platelets (Cloutier et al., 2018). This surge in serum serotonin could lead to endogenous activation of BJR, as bolus intravenous infusion of serotonin reliably triggers BJR experimentally(Fozard, 1983; Whalen et al., 2000). Additional seizure related disturbances like central or obstructive apneas may contribute to BJR activation. Hypoxia, which could result from ictal apnea, is known to increase excitatory neurotransmission to cardiac vagal motor neurons that cause vagal bradycardia (Griffioen et al., 2007) and induces platelet activation (Tyagi et al., 2014) which is the main source of circulating serotonin. As such, hypoxia resulting from apnea may increase likelihood of exaggerated BJR during seizures. Consistent with this, individual SUDEP case reports indicate increases in parasympathetic activity including triggered bradycardia during terminal SUDEP events (Jeppesen et al., 2019), providing further evidence that activation of the parasympathetic-mediated bradycardia during the BJR is clinically relevant. This increased BJR responsivity could also trigger the exaggerated ‘low pressure’ baroreflex via resulting hypotension thereby further complicating homeostasis. Therefore, the magnitude and timing of increased BJR in *Kcc2/Crh* mice suggests a critical role in the generation of robust bradycardia during spontaneous seizures that could link to sudden death.

Chronic inhibition of vagal parasympathetic motor output (the driver of BJR reflex bradycardia) improved mortality by 10% in Kcc2/Crh mice. Although this improvement provides some hope for patients at high risk for SUDEP with no treatment options, additional avenues of investigation are needed to more directly link seizure-related bradycardias to vagal parasympathetic motor output. In addition to bradycardia, the BJR induces a significant apnea and depressor response. Since SUDEP events include all of these physiological responses, it remains possible that elimination of one can only prevent mortality in certain incidences or in specific individuals. Similar to models proposed for other forms of sudden death (e.g. sudden infant death syndrome) (Kinney et al., 2009), SUDEP could result from “a perfect storm” of multiplicative homeostatic risk factors. As the BJR results in a depressor response, if sufficiently severe, it could trigger “low pressure” baroreflex-mediated paradoxical bradycardia. Unlike BJR-mediated bradycardia, this “low pressure” baroreflex-mediated paradoxical bradycardic reflex response is mediated by sympathetic reflex circuits (Souza et al., 2022) and would not be blocked by the scopolamine used here. Results here suggest the bradycardia from 5HT3R activation alone is sufficient to trigger SUDEP in ∼10% of *Kcc2/Crh* mice. However, the apnea and depressor response may either contribute directly or serve as triggers for secondary reflex responses to promote SUDEP. BJR reflex responsivity could serve as a biomarker for patients at highest risk for SUDEP; patients who would benefit from increased vigilance in monitoring. Future directions could further examine how reflex circuits are modulated singly, and in combination, and whether they could serve as biomarkers in SUDEP.

### Future Directions

Although the present study includes only non-fatal seizures, our report of HR during spontaneous seizure events supports work suggesting physiological events during non-fatal seizures predict SUDEP risk (Lamrani et al., 2023; Ryvlin, Nashef, & Tomson, 2013; Schuele et al., 2011). It remains to be determined if ictal events during fatal and non-fatal spontaneous seizures are different and future studies could help clarify any distinctions. Our examination focused on HR (and not respiration or BP). Whether exaggerated BJR-mediated HR response co-occurs with greater magnitude BJR-mediated apnea and hypotension remains to be determined. Finally, the cascade of stress signaling and cardiorespiratory disturbances examined here could have forms of state-dependency during both ictal and interictal periods.

The precise functional role of potential BJR bradycardia during seizures warrants additional studies. A recent report suggested that in addition to the classic triad of physiological responses, BJR activation transiently decreases EEG power (Lovelace et al., 2023). Initiation of ictal asystole associates with early seizure termination in patients (Moseley et al., 2011; Schuele et al., 2010) and suppression of cerebral electrical activity (Rossetti et al., 2005). We observed the most prominent bradycardia at the end of seizures (Fig. 2). It is possible BJR activation during seizures promotes seizure termination and/or post-ictal EEG suppression. However, continued activation and strengthening of the BJR could ultimately prove detrimental. A role for excessive BJR activation to help facilitate seizure control and arousal, that leads to negative plasticity, is consistent with the “two-hit” pathomechanism theory of SUDEP (Mueller et al., 2018).

### Conclusions

TLE increased susceptibility to SUDEP in Kcc2/Crh mice and induced significant alternations in their cardiac function, namely an early resting tachycardia and modification of baroreflex parameters. TLE in the context of hyperreactive PVN^Crh^ neurons, increases abnormal cardiac homeostatic regulation by exaggerating BJR-mediated bradycardia and “low pressure” baroreflex-mediated paradoxical bradycardia. These increased bradycardiac responses are suited triggers for increased severity of ictal bradycardia seen in TLE mice with increased stress signaling (*Kcc2/Crh*). Ictal activation of these improperly weighted reflexes could result in the cardiorespiratory collapse seen during SUDEP. Together, results here provide novel mechanisms of SUDEP in the context of hyperreactive stress central circuits. Future research can expand whether preventing activation of one or more autonomic reflexes during seizures can decrease SUDEP susceptibility.

## MATERIALS AND METHODS

### Animals

All animal procedures were performed in accordance with NIH standards, ARRIVE guidelines for the care and use of laboratory animals, and approval by the University of Missouri Animal Care and Use Committee (ACUC). *Kcc2^f/f^* (WT) and *Kcc2/Crh* mice (both sexes; 8-35 weeks of age) were initially acquired as a gift in kind (Dr. Jamie Maguire)(Basu et al., 2024; Melon et al., 2018) to establish a live colony. Animals had *ad libitum* access to food and water and were housed under 12h light – 12h dark conditions.

### Telemetry implantation for recordings of electroencephalogram (EEG) and electrocardiogram (ECG)

Mice were implanted with HD-X02 (Data Science International) telemetry devices to examine changes in resting HR (via electrocardiogram, ECG), as well as changes in HR during spontaneous seizures (recorded with electroencephalogram, EEG) over time. Mice were anesthetized with isoflurane in oxygen (4% induction, then maintained at 2%) to effect (lack of toe pinch). Surgical sites were aseptically prepared, and mice placed on a heating pad for supplemental heat. A transverse incision was made through the skin on the dorsal flank. Using blunt tipped scissors, a subcutaneous pocket was created and the implant inserted. One set of bipotential leads was tunneled to two ventral incision sites and placed in a Lead II ECG configuration. Leads were secured in place and incision site closed with 5–0 silk suture. Mice were then placed in a stereotaxic apparatus in sternal recumbency. An incision was made along the dorsal midline and bipotential leads were tunneled to the incision in the scalp. Two burr holes were made into the skull (one ∼1mm anterior to bregma and ∼1mm to the right of midline and the other at ∼2 mm posterior to bregma and ∼1mm to the left of midline). EEG screws (0-80 × 1/16, PlasticsOne, Roanoke, VA) were then inserted and secured to the skull with biopotential leads attached. Dental acrylic was applied to cover screws. Incisions were closed with either 5-0 silk suture or wound clips (FST). Mice were given subcutaneous pain medication (Buprenorphine extended release, Ethiqa XR, 1mg/kg) and placed back in their home cage with supplemental heat and monitored postoperatively. Wound clips/sutures were removed 10-14 days after surgery.

### Seizure induction via ventral intrahippocampal kainic acid microinjection

The ventral intrahippocampal kainic acid (vIHKA) model, an established model of temporal lobe Epilepsy, was used to generate chronically epileptic mice as previously described (Basu et al., 2024; Zeidler et al., 2018). Briefly, mice were anesthetized with a ketamine/xylazine cocktail (100mg/kg ketamine; 10mg/kg xylazine) and aseptically prepared for stereotaxic surgery as detailed above. A burr hole was made at established coordinates (ventral hippocampus; -3.6mm from bregma, 2.8mm from midline)(Basu et al., 2024; Paxinos & Franklin, 2001). A 34 gauge needle (WPI) attached to a 10uL Hamilton syringe was then lowered into the ventral hippocampus (-2.8mm from surface) and kainic acid (KA; Millipore Sigma, 20mM, 100nL) or sterile saline (100nL; 0.9%) was microinjected using a microsyringe pump (UltraMicroPump3 and SMARTouch Controller, WPI) at a rate of 1.7ul/sec. The needle was then slowly withdrawn. Incisions were closed with VetBond (3M) tissue adhesive or 5-0 silk suture. Mice were given warmed subcutaneous fluids (1 mL, 0.9% Saline) and pain medication (Buprenorphine extended release, Ethiqa XR, 1mg/kg) and placed back in their home cage with supplemental heat and monitored twice daily for duration of study. Seizures were not pharmacologically terminated, and *status epilepticus* was observed upon waking.

### ECG and EEG Telemetry

Mice were allowed to recover from the telemetry implant surgery and then acclimated for at least three days prior to telemetry recordings. Animals were then recorded for one day (at least a week after telemetry implant surgery) to obtain baseline measurements (prior to vIHKA or saline hippocampal injections). This recording occurred between the hours of 10:00 and 17:00 h in all cohorts, and animals were singly housed during this time with water and food available *ab libitum*.

After vIHKA or saline hippocampal injection, telemeter implanted mice were recorded weekly (day 7, 14, 21, 28) for ECG and EEG. To note, two WT mice (1 saline hippocampal injected, 1 vIHKA) were not recorded on day 28 due to issues with the telemeters, not SUDEP. Signals from telemetry probes were acquired at a sampling frequency of 1 kHz using the DSITalker interface (Cambridge Electronic Design Limited) connected to MX2 PhysioTel telemetry hardware (Data Sciences Inc.)

### Autonomic reflex testing

All autonomic reflex testing was performed using a jugular catheter tunneled into the right atria as previously detailed (Strain et al., 2023). Briefly, a jugular catheter was placed under urethane anesthesia (1500 mg/kg w/supplemental doses as needed) immediately before autonomic reflex testing. Surgical sites were aseptically prepared. A midline incision and blunt dissection was completed to visualize the right external jugular vein with a surgical microscope. The jugular vein was gently lifted with curved forceps, and two small sutures placed underneath. The cranial suture was used to ligate the jugular vein. Using fine spring scissors, a small incision was made into the jugular vein and a beveled catheter (polyethylene tubing; PE10; Intramedic, 427401) containing saline was inserted and secured with suture.

#### For Baroreflex

In experiments where baroreflex was assessed, blood pressure (BP) and HR were recorded using HDX11 telemetry probes. Briefly, a transverse incision was made through the skin on the dorsal flank. A subcutaneous pocket was created via blunt dissection. Bipotential leads and catheter were tunneled, using a trocar, to the ventral incision site on the neck. The ECG leads were further tunneled and secured in a Lead II ECG configuration. The carotid artery was isolated using blunt force dissection and three sutures placed underneath the artery. The cranial suture was ligated to the carotid artery just proximal to the carotid bifurcation. To allow placement of the catheter, the most distal suture was used to temporarily occlude blood flow. Using the 25-gauge needle, the artery was pierced, and catheter inserted. The middle suture was secured and occlusion suture released. The catheter was then advanced towards the aortic arch. The middle and occlusion suture were tightened and incisions closed with suture. Sodium nitroprusside (SNP, 600ug/kg) was infused (0.04 mL/min; 1mg/mL) into the jugular vein to decrease BP. Phenylephrine (PE, 15mg/kg) was infused (0.04 mL/min, 25mg/mL)(Karlen-Amarante et al., 2012) to increase BP.

#### For Bezold Jarisch Reflex (BJR)

In experiments where the BJR was assessed, HR was recorded either via previously implanted HDX02 probe (animals that underwent chronic telemetry recordings) or needle electrodes inserted into each forepaw in a lead I ECG configuration. ECG recordings were AC amplified (1000x; CED 1902 amplifier, Cambridge Electronic Design, Cambridge, UK), band pass filtered (1-1000Hz) and digitized at 1 kHz (CED Power 1401, Cambridge Electronic Design, Cambridge, UK). Phenylbiguanide (50ug/kg), a serotonin type 3 receptor (5HT3R) agonist, was infused (1mL/min, 0.1mg/mL) via the jugular catheter. In a subset of mice, animals were tested to ensure autonomic contributions by subsequent administration of the sympathetic antagonist, atenolol (10 mg/kg, I.P.) and then parasympathetic antagonist, scopolamine (1 mg/kg, I.P), with PBG administration following each drug. At least 15 minutes was allowed between repeated testing to eliminate tachyphylaxis and allow for full antagonist effects.

### Chronic muscarinic blockade

In a subset of vIHKA-injected *Kcc2/Crh* mice, an osmotic mini pump (Alzet Model 1004) containing methyscopolamine was implanted four days after seizure induction to test the capacity for chronic parasympathetic blockade to prevent SUDEP mortality. Briefly, mice were anesthetized with isoflurane in oxygen (4% induction, then maintained at 2%) to effect (lack of toe pinch) and aseptically prepared for implantation as detailed for telemeters. A transverse incision was made through the skin on the dorsal flank. Using blunt tipped scissors, a subcutaneous pocket was created. The pocket was then irrigated with sterile saline and a mini pump containing methylscopolamine (12mg/kg/day, n=10) was inserted. The site was then closed with wound clips (FST). Mice were given pain medication cocktail (buprenorphine 0.1 mg/kg and carprofen 10 mg/kg) and placed back in their home cage with supplemental heat and monitored postoperatively. Wound clips were removed a week after surgery.

### Immunohistochemistry

For immunohistochemical analysis, mice were deeply anesthetized with isoflurane or urethane (if following reflex testing) until the toe-pinch reflex was absent. Mice were decapitated and brains removed and bisected down the midline. Brains were fixed by submerging in 2.5% PFA for 24 hours and then cryoprotected in 20% sucrose for 24 hours followed by 30% sucrose for 24 hours. Hippocampal sections were cut at 30 µm in the coronal plane with a cryostat (Leica CM1860). Serial sections through the ventral hippocampus were rinsed with PBS (pH 7.4). Nonspecific immunoreactivity was blocked with 5% normal donkey serum (Jackson Immunoresearch, ref#017-000-121) in PBS + Triton (0.3%). ZnT3 was identified with primary rabbit anti-Znt3(1:100, Synaptic Systems) followed by secondary donkey anti-rabbit 488 Alexa Fluor (1:200, Invitrogen). Slices were mounted using DAPI mounting media. Tissue from all groups were run in parallel with the same antibody cocktails. Images were acquired under identical parameters, including exposure time and illumination intensity. Further processing of brightness and contrast were performed identically for figure presentation. Negative controls were run without primary antibody. All imaging was done with a fluorescence microscope (BZ-X800 Keyence, Itasca, Illinois) using filters appropriate for the two fluorescent dyes.

### Drugs

(-)Scopolamine methyl bromide, (R)-(-)- Phenylephrine hydrochloride, Atenolol, and Sodium nitroprusside dihydrate were from Sigma-Aldrich (St Louis, MO, USA). 1-Phenylbiguanide hydrochloride was from Combi-Blocks (San Diego, CA, USA).

### Data analysis

For analysis of mortality, only animals that were taken out to endpoint (28 days) or were found dead in their cage (presumed SUDEP) were included. Animals were checked twice daily for mortality.

All HR recordings were analyzed offline in Spike2 software. Resting HR was averaged from a stable two-hour recording period at least two hours after the animals were placed in the recording room. Collected EEG data was analyzed using SeizyML for spontaneous seizure onset and termination (Antonoudiou et al., 2025). Time from seizure onset to termination was used to determine seizure duration. For analysis of changes in HR during seizures, a significant change in HR was considered if HR changed (increased or decreased) by > 5% of HR during the seizure. As a tachycardia often occurred before the onset of the seizure, the baseline period was sampled for 60 secs during the preictal period. For analysis of postictal HR, HR was averaged for the first 10 seconds immediately following seizure termination and then the following 20 seconds. For qualitative representation of the grouped average HR during seizures, a stimulation histogram was generated in Spike2 triggered by seizure termination. Beats were binned into 1 sec intervals.

For analysis of BJR, HR was averaged 30 sec prior to phenylbiguanide infusion and then again in two, two second bins. The minimum of the first two bins was selected to represent BJR magnitude. For analysis of HR and BP during sodium nitroprusside infusion, BP and HR were averaged for one minute before infusion to represent baseline. HR and BP were then averaged for 10 sec at the end of the infusion to represent the “early” response. Right before PE administration (∼30-45 sec after SNP infusion), HR and BP were averaged for 10 secs to represent the “late” response. Both early and late BP and HR values were subtracted from baseline values to capture the biphasic HR response to SNP. To analyze HR and BP changes with PE administration, binned averages (0.5 sec duration) of BP and HR were sampled over time during PE infusion. Change in HR by Change in BP was then fitted with a modified WYSIWYG Boltzmann sigmoidal equation (Heusser et al., 2021) which generated V50 (as an index of BP set point) and slope (as an index of baroreflex sensitivity) values. The change in BP with PE administration was calculated as the difference before PE infusion and an average BP over 10 sec at the peak of the PE-induced BP response. For terminal reflex testing and/or in vivo telemetry, animals with baseline HR and/or BP two standard deviations above or below means seen in control groups were excluded from analysis.

For immunohistochemical analysis, ventral hippocampus was identified with the use of a mouse brain stereotaxic atlas and DAPI immunoreactivity (Paxinos & Franklin, 2001). Mossy fiber sprouting and dentate granule cell density were quantified in ImageJ (version 1.54f). For each slice, the entire granule cell layer and molecular layer of the dentate gyrus were labeled as a region of interest (ROI) and mean intensity of the ROI was calculated. Mean fluorescence intensity of the ROI was averaged across two sections to quantify mossy fiber sprouting for each animal. The cell counter function in ImageJ was used to manually count cells (DAPI+) in a region (containing at least 50 cells) of the dentate gyrus in the upper blade, with a ROI around the counted cells used to determine area. Cell density was quantified as number of cells per area. The experimenter was blinded to treatment during analysis.

### Statistics

Data are presented as mean ± s.d. unless otherwise stated or shown as individual points and means to highlight individual responses. When two groups of independent samples were compared, a two-tailed unpaired t-test was used. In experiments with one main effect, a one-way ANOVA, with or without repeated measures (when appropriate), was used to test for the main effect. One-way ANOVA was followed up with Tukey’s multiple comparisons test. In experiments with two main effects, a two-way ANOVA, with or without repeated measures (when appropriate), was used to test the two main effects and the interaction effect. Two-way ANOVA was followed up with Tukey’s multiple comparisons test unless otherwise specified. To analyze repeated-measures data with missing values (HR over time w/SUDEP mortality), a mixed effects model was used and followed up with Tukey’s multiple comparisons test. Significance was accepted when P < 0.05. All analyses were performed using GraphPad Prism (v10.5.0, San Diego, CA, USA).

## Authors contributions

S.E.S., J.A.B, J.L.M, and C.R.B conceived and designed research; S.E.S., K.E.D, and G.E.B. performed experiments; S.E.S. and C.R.B analyzed data; S.E.S. and C.R.B. interpreted results of experiments; S.E.S. prepared figures; S.E.S. drafted manuscript; S.E.S., C.R.B, J.A.B, and J.L.M edited and revised manuscript; S.E.S., K.E.D, G.E.B, J.A.B, J.L.M, and C.R.B. approved final version of manuscript.

## Supplement

**Table 1.1:**
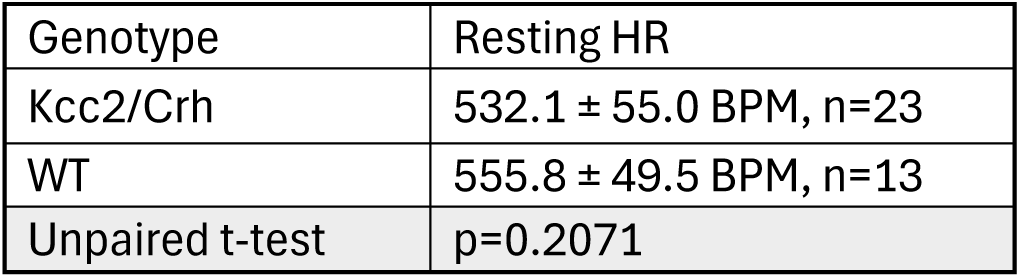
Figure 1A

**Table 1.2:**
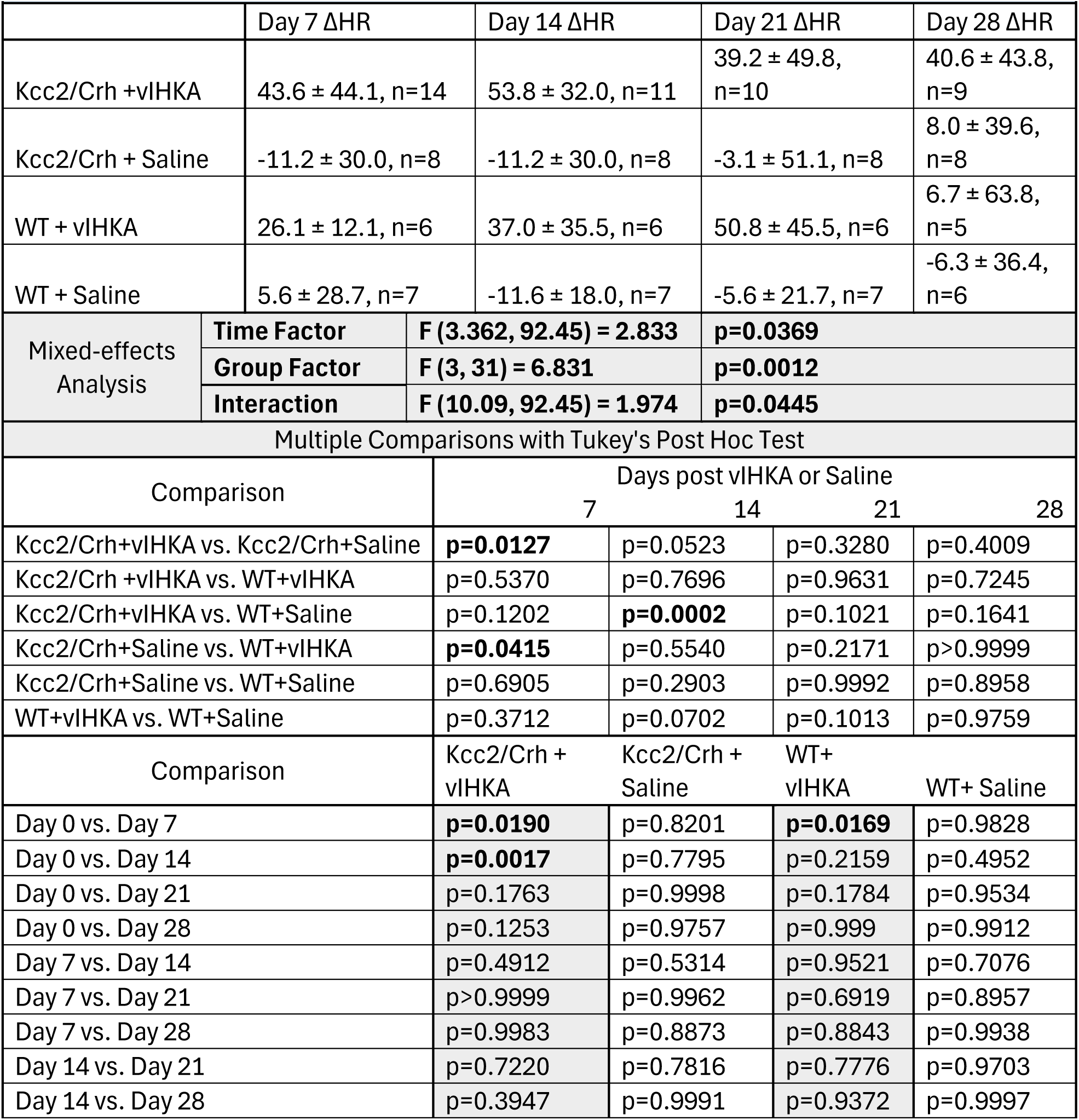
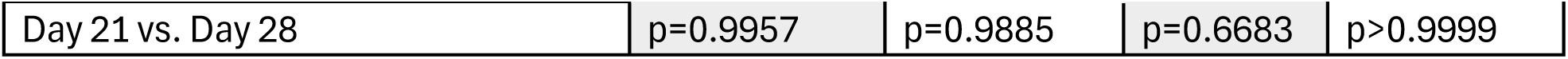
Figure 1C- HR normalized to Day 0 (BPM)

**Table 2.1:**
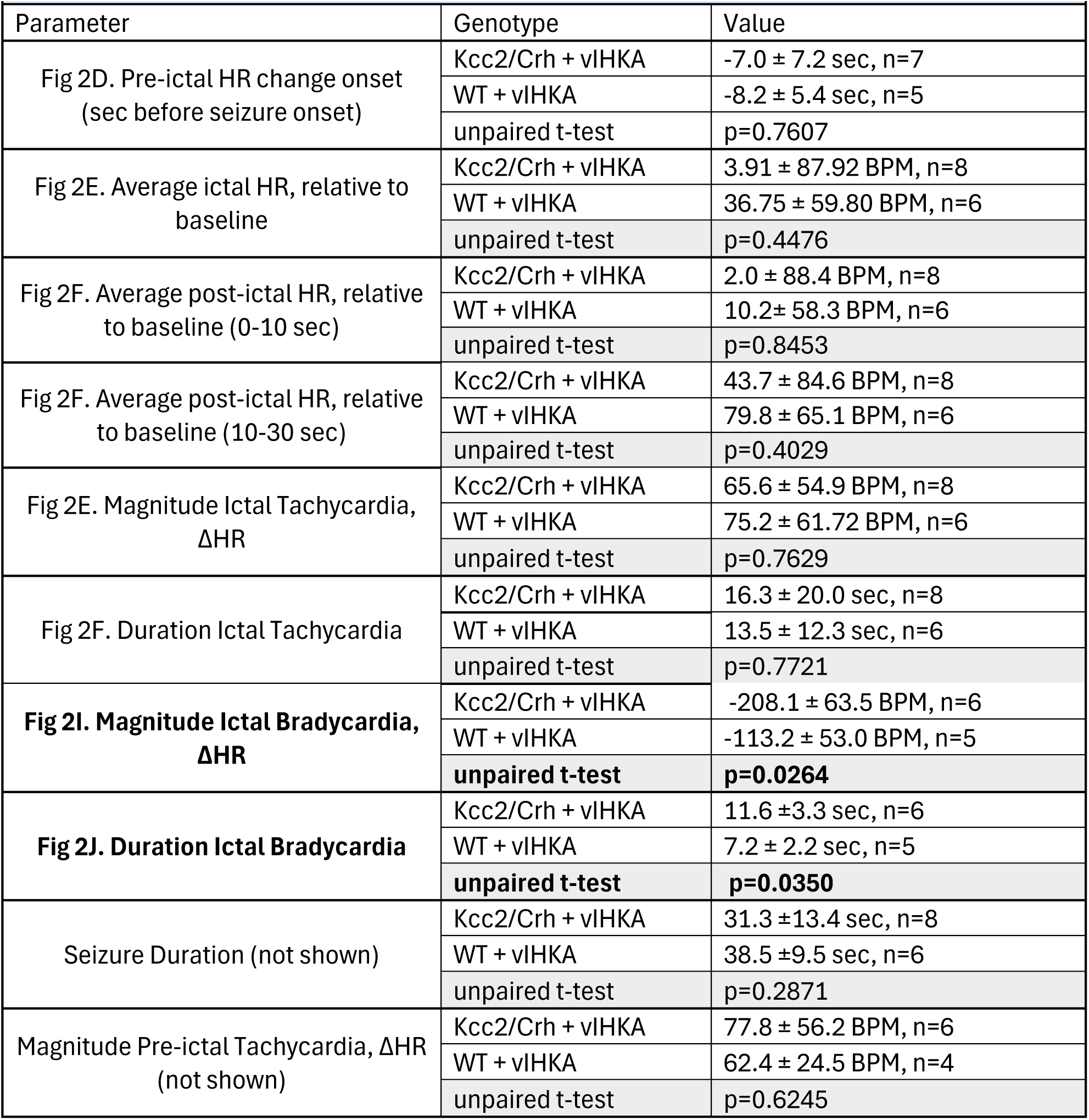
Figure 2

**Table 3.1:**
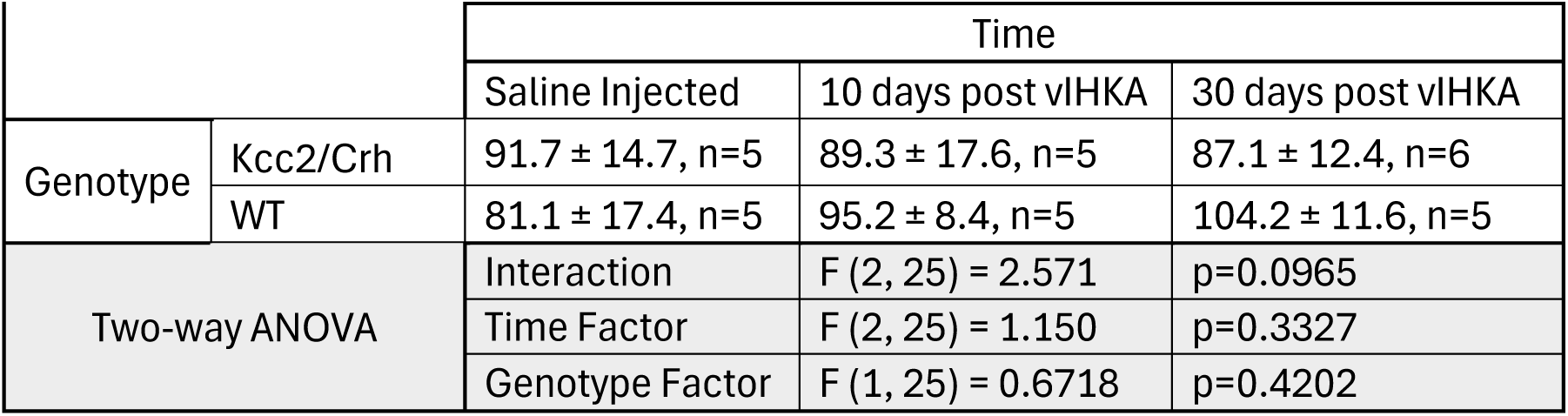
Figure 3C- Phenylephrine mediated increase in BP (ΔmmHg)

**Table 3.2:**
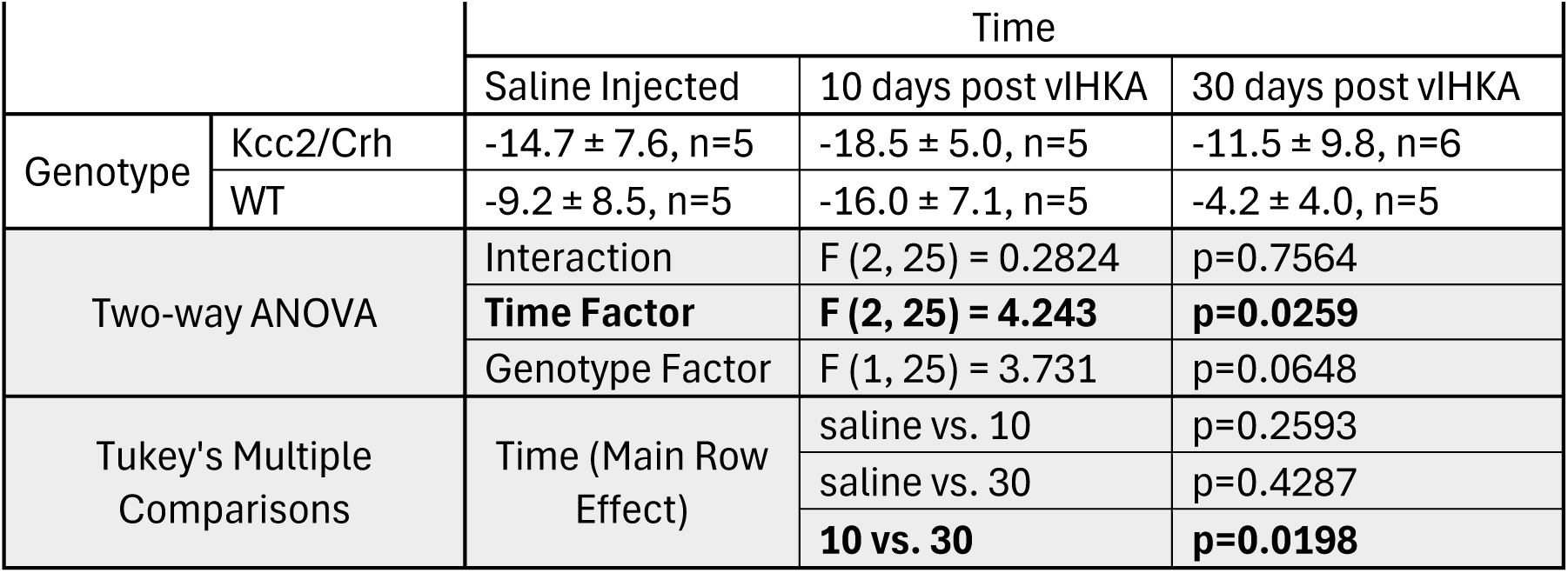
Figure 3E- Slope (ΔBPM/ΔmmHg)

**Table 3.3:**
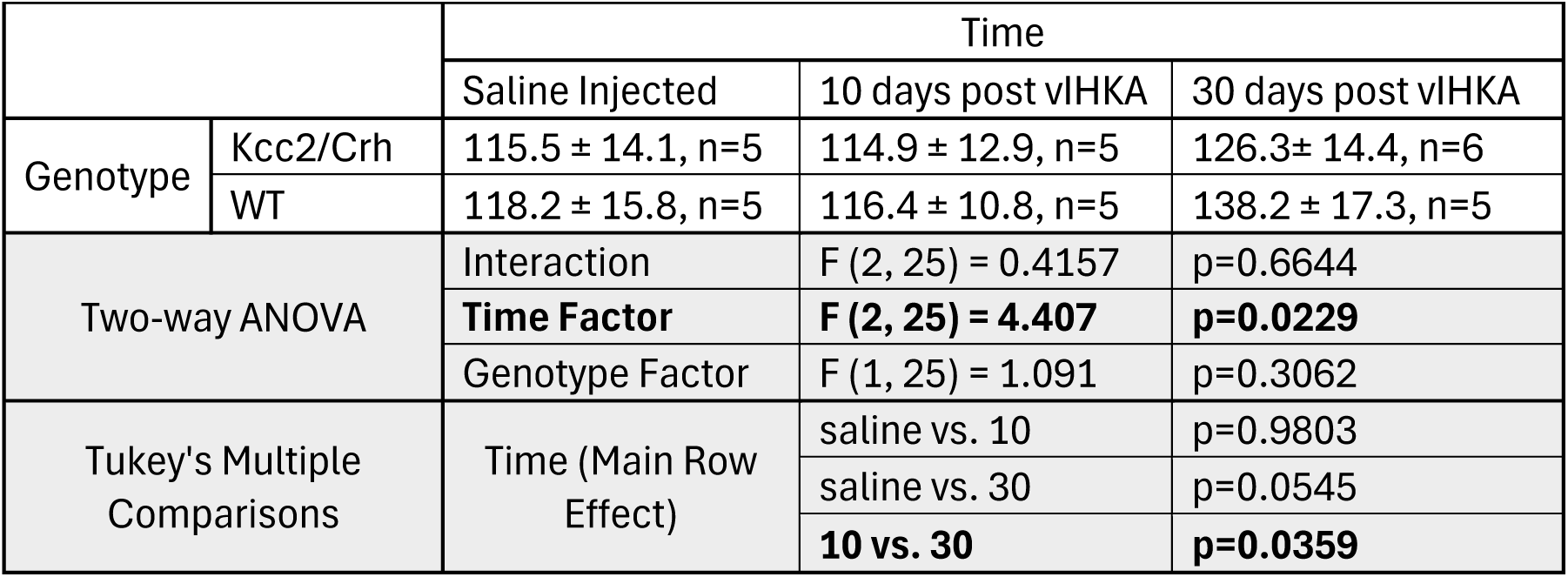
Figure 3D- V50 (mmHg)

**Table 4.1:**
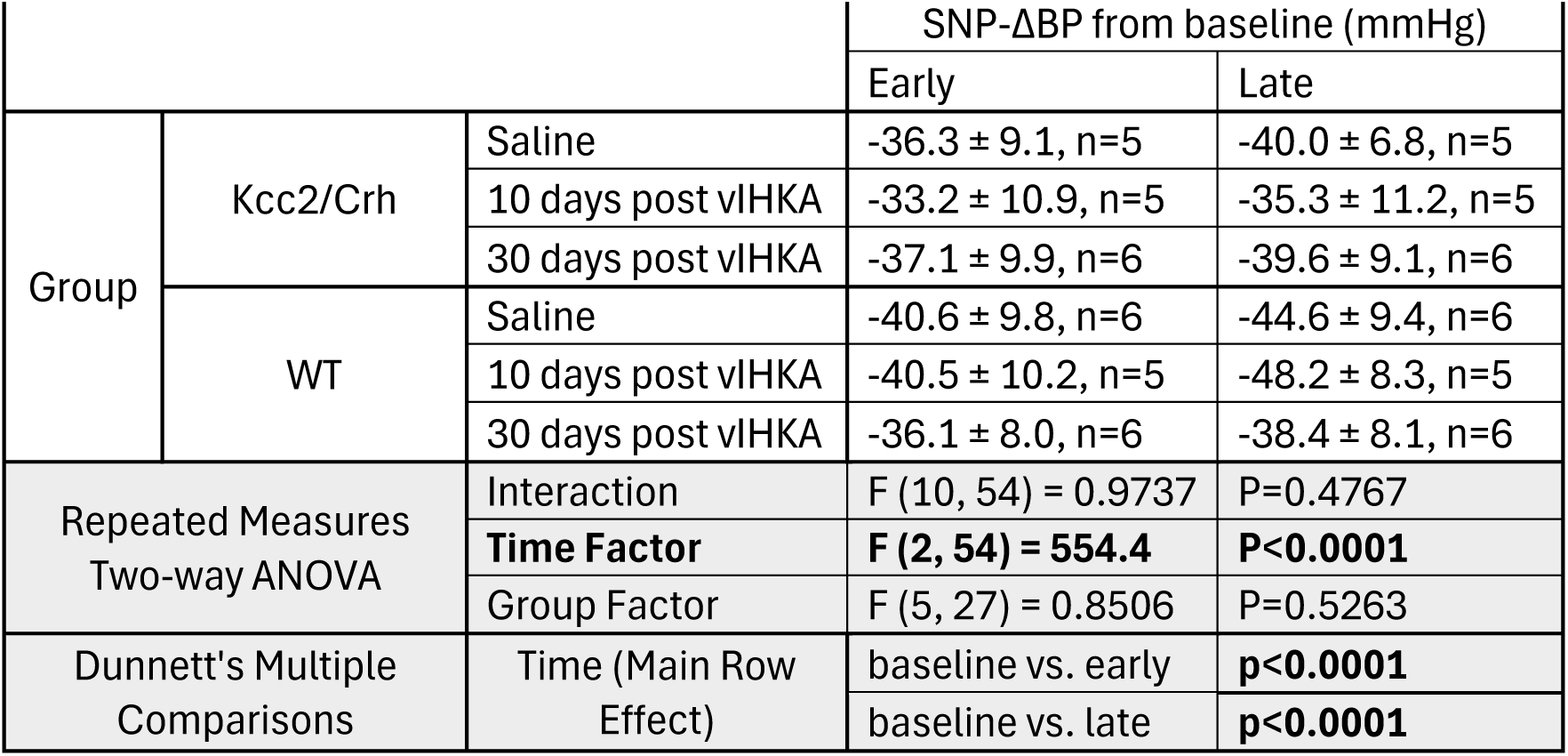
Figure 4C

**Table 4.2:**
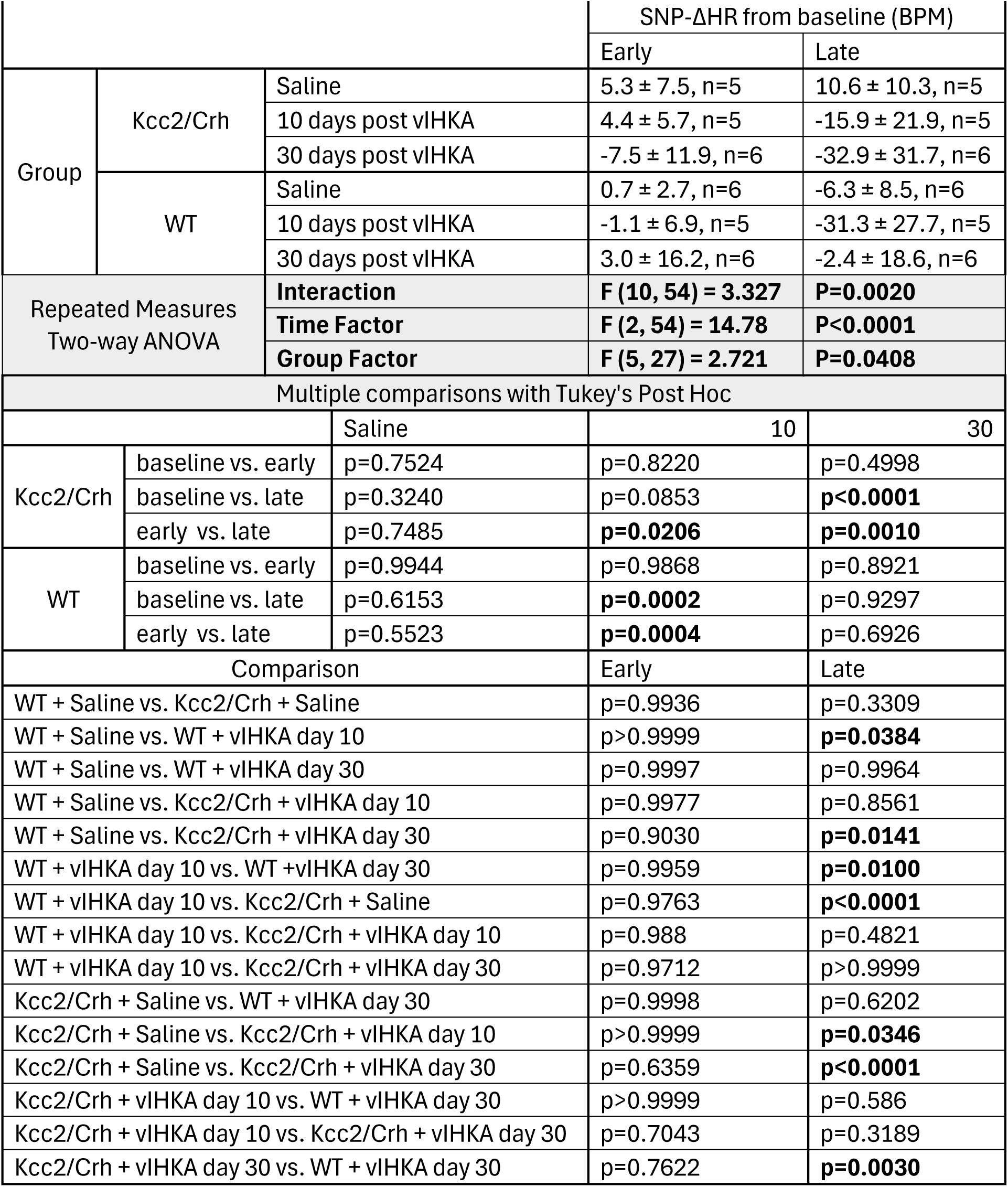
Figure 4D

**Table 4.3:**
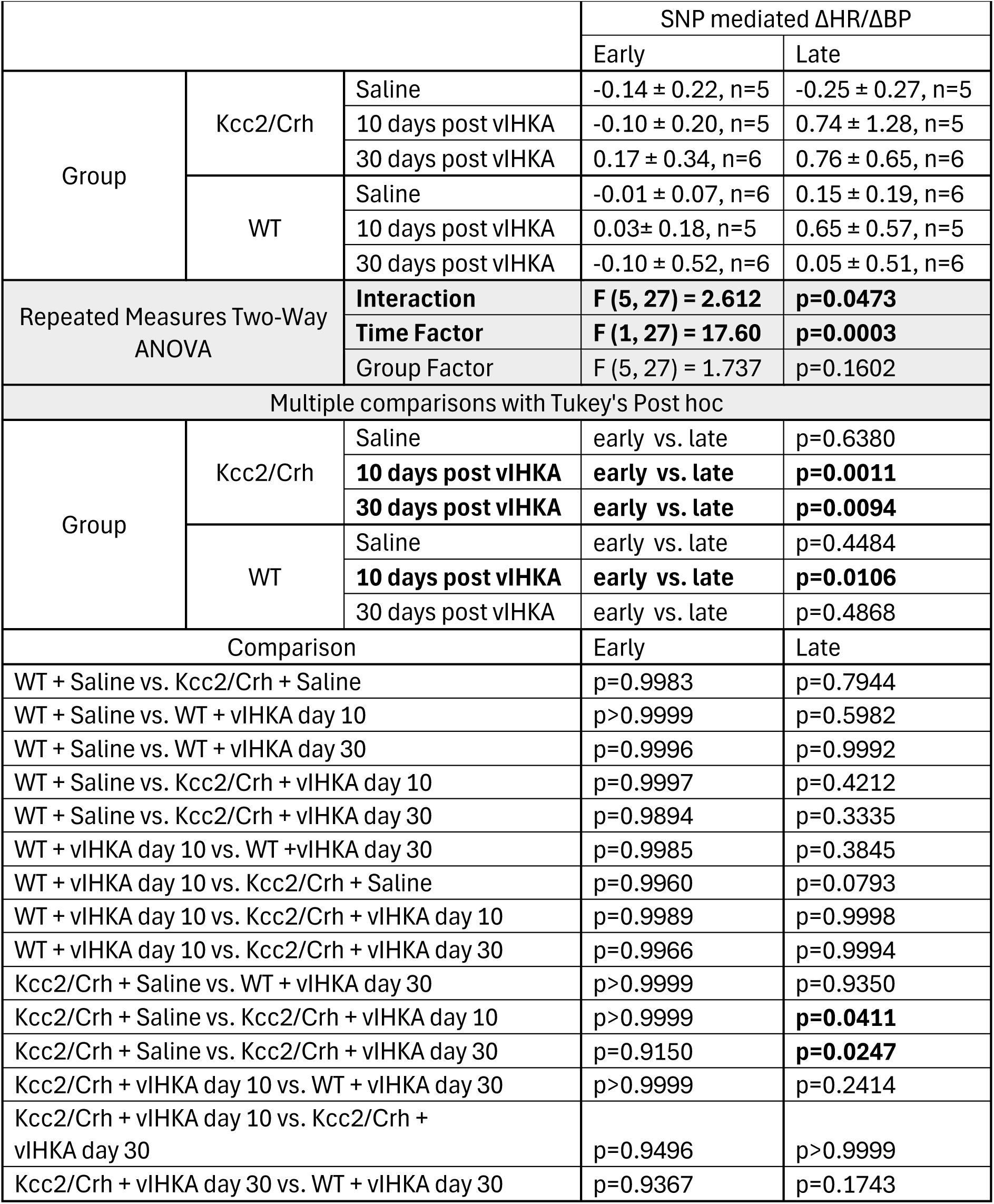
Figure 4E

**Table 5.1:**
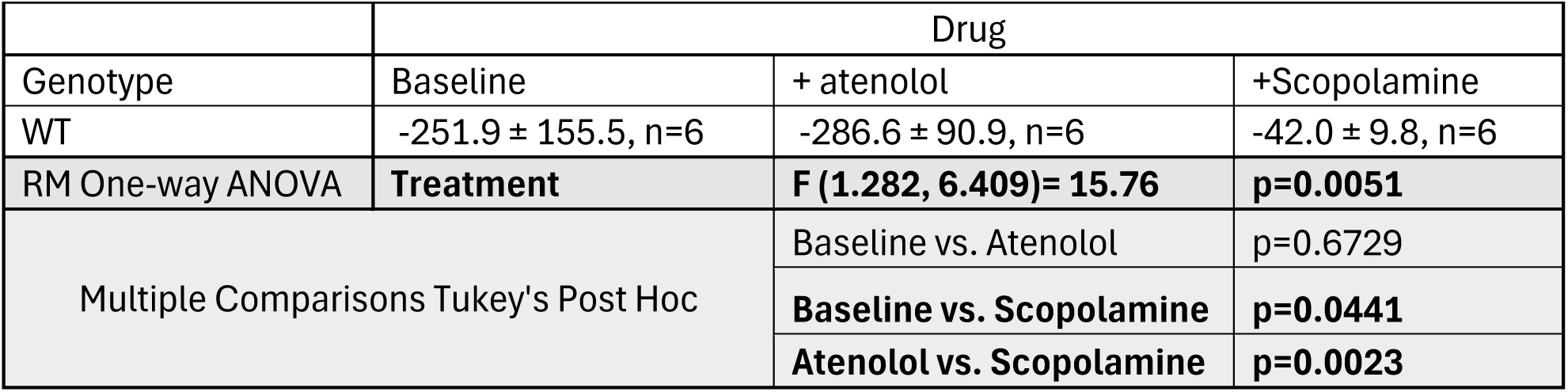
Figure 5B- PBG mediated ΔHR (BPM)

| Table 5.1: Figure 5B- PBG mediated ΔHR (BPM) | | | |
| --- | --- | --- | --- |
|  | Drug |  |  |
| Genotype | Baseline | + atenolol | +Scopolamine |
| WT | -251.9 ± 155.5, n=6 | -286.6 ± 90.9, n=6 | -42.0 ± 9.8, n=6 |
| RM One-way ANOVA | **Treatment** | **F (1.282, 6.409)= 15.76** | **p=0.0051** |
| Multiple Comparisons Tukey's Post Hoc |  | Baseline vs. Atenolol | p=0.6729 |
|  |  | **Baseline vs. Scopolamine** | **p=0.0441** |
|  |  | **Atenolol vs. Scopolamine** | **p=0.0023** |

**Table 5.2:**
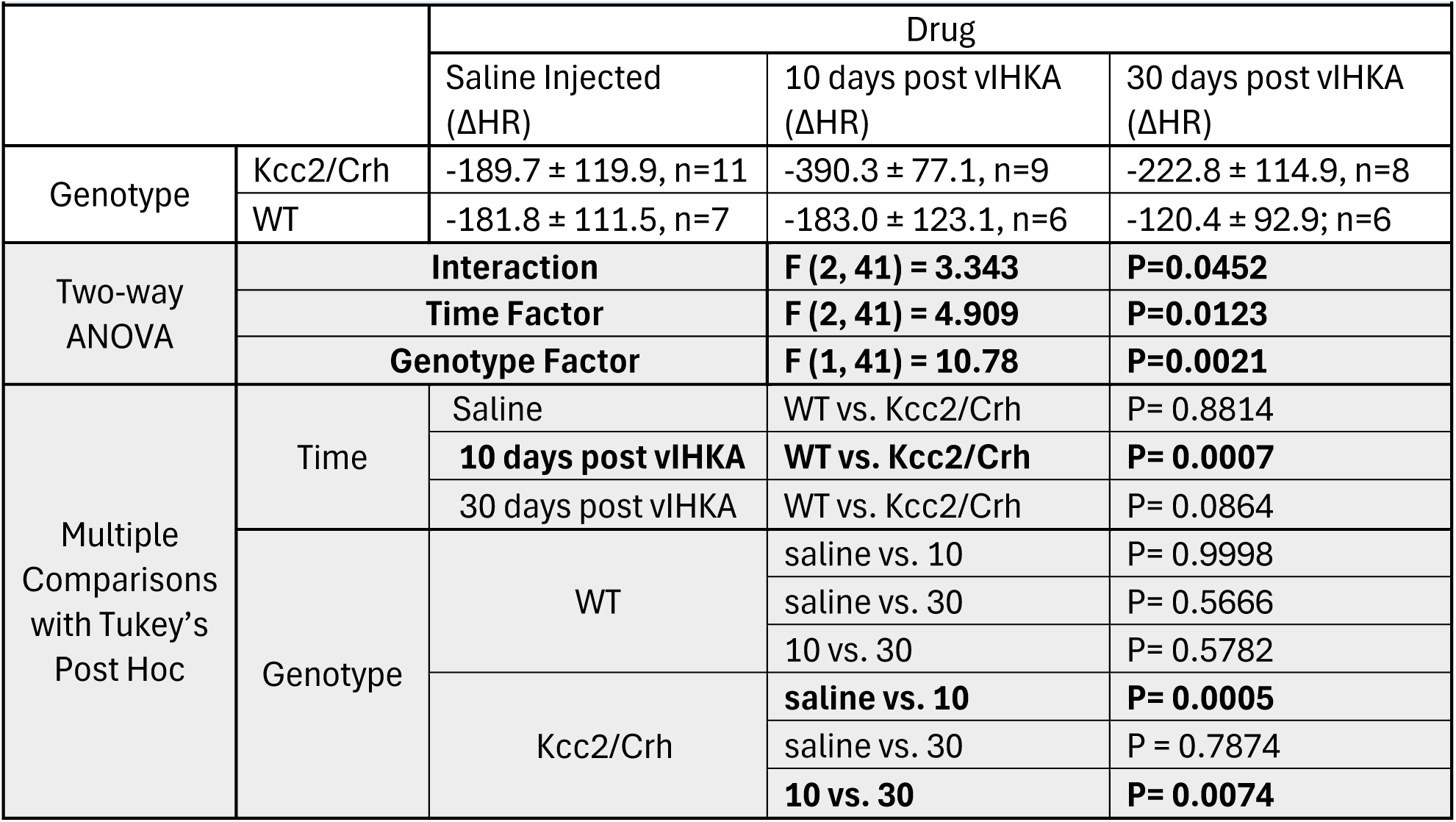
Figure 5E- PBG mediated ΔHR over time (BPM)

**Table 6.1:**
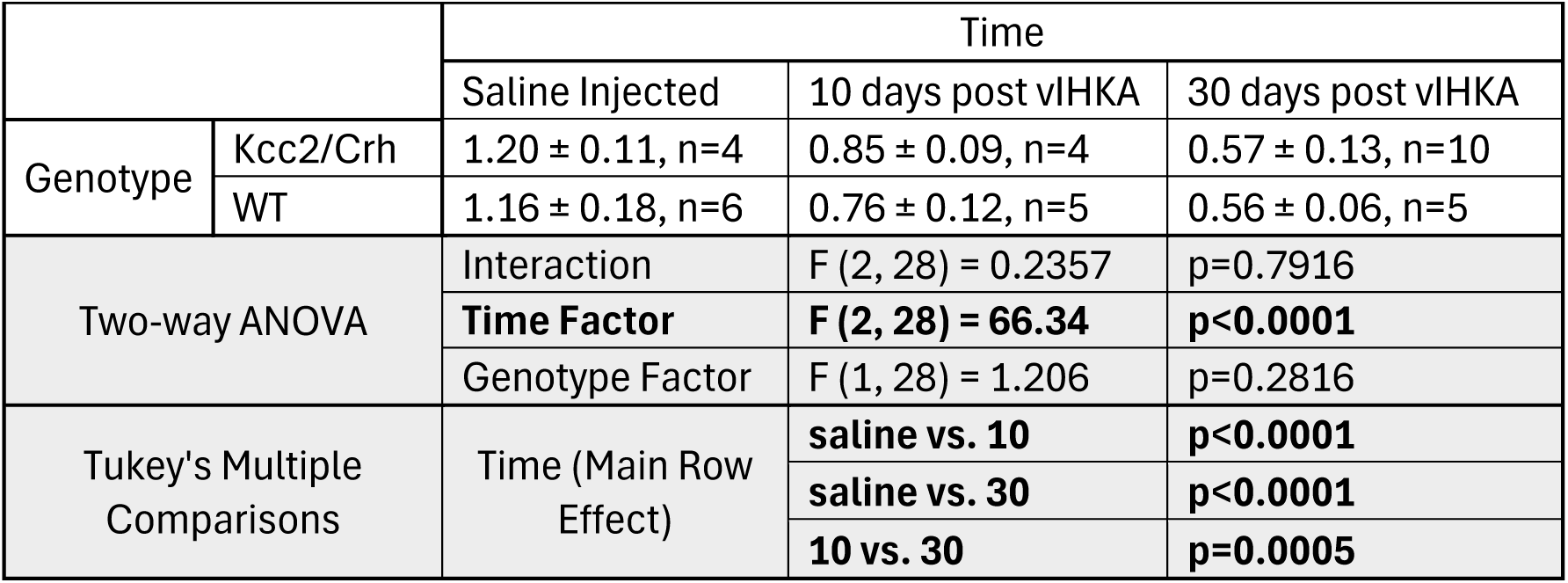
Figure 6B- Dentate Granule Cell Density (# cells/area)

**Table 6.2:**
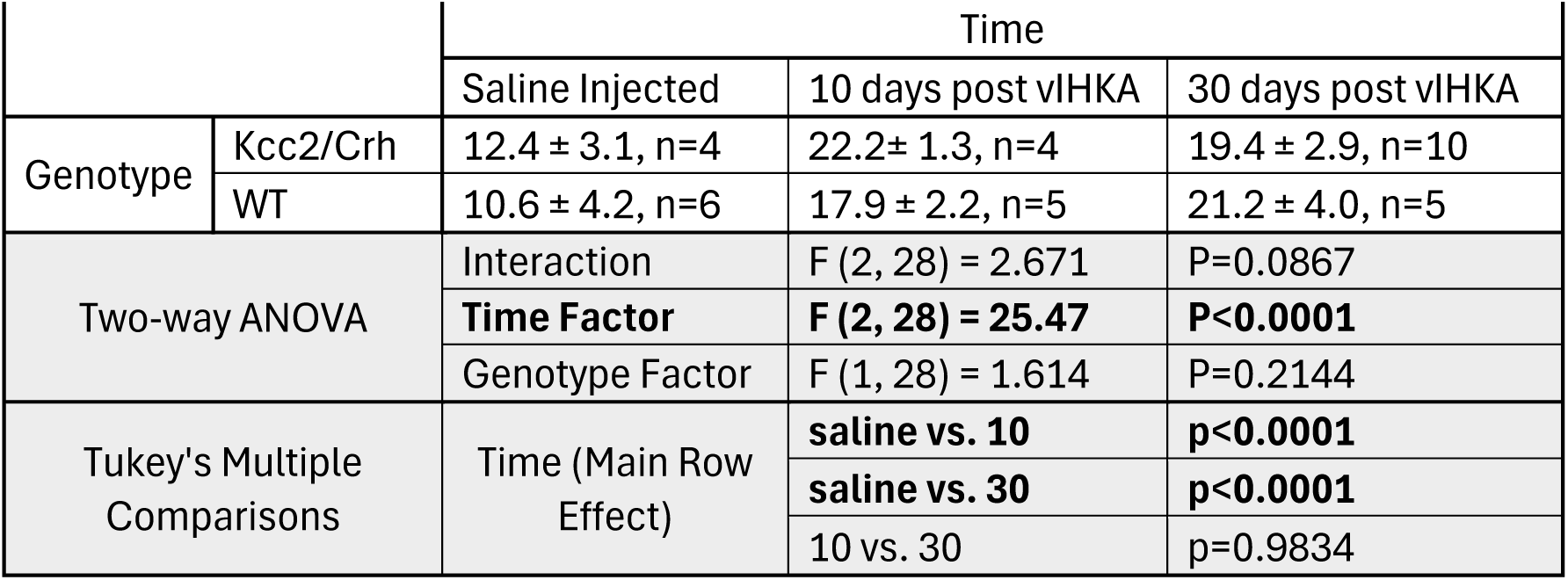
Figure 6C- Mossy Fiber Sprouting (Mean Fluorescence Intensity)

## References

Antonoudiou, P., Basu, T., & Maguire, J. (2025). SeizyML: An Application for Semi-Automated Seizure Detection Using Interpretable Machine Learning Models. Neuroinformatics, 23(2), 23. 10.1007/s12021-025-09719-4

Basu, T., Antonoudiou, P., Weiss, G. L., Coleman, E. M., David, J., Friedman, D., Laze, J., Strain, M. M., Devinsky, O., Boychuk, C. R., & Maguire, J. (2024). Hypothalamic-Pituitary-Adrenal Axis Dysfunction Elevates SUDEP Risk in a Sex-Specific Manner. eNeuro, 11(7). 10.1523/ENEURO.0162-24.2024

Bozorgi, A., Chung, S., Kaffashi, F., Loparo, K. A., Sahoo, S., Zhang, G. Q., Kaiboriboon, K., & Lhatoo, S. D. (2013). Significant postictal hypotension: expanding the spectrum of seizure-induced autonomic dysregulation. Epilepsia, 54(9), e127–130. 10.1111/epi.12251

Britton, J. W., Ghearing, G. R., Benarroch, E. E., & Cascino, G. D. (2006). The ictal bradycardia syndrome: localization and lateralization. Epilepsia, 47(4), 737–744. 10.1111/j.1528-1167.2006.00509.x

Budde, R. B., Pederson, D. J., Biggs, E. N., Jefferys, J. G. R., & Irazoqui, P. P. (2020). Mechanisms and prevention of acid reflux induced laryngospasm in seizing rats. Epilepsy Behav, 111, 107188. 10.1016/j.yebeh.2020.107188

Cloutier, N., Allaeys, I., Marcoux, G., Machlus, K. R., Mailhot, B., Zufferey, A., Levesque, T., Becker, Y., Tessandier, N., Melki, I., Zhi, H., Poirier, G., Rondina, M. T., Italiano, J. E., Flamand, L., McKenzie, S. E., Cote, F., Nieswandt, B., Khan, W. I.,…Boilard, E. (2018). Platelets release pathogenic serotonin and return to circulation after immune complex-mediated sequestration. Proc Natl Acad Sci U S A, 115(7), E1550–E1559. 10.1073/pnas.1720553115

Cramer, W. (1915). ON THE ACTION OF VERATRUM VIRIDE: WITH SOME REMARKS ON THE INTERRELATIONSHIP OF THE MEDULLARY CENTRES. The Journal of Pharmacology and Experimental Therapeutics, 7(1), 63–82.

Demetriades, D., Chan, L. S., Bhasin, P., Berne, T. V., Ramicone, E., Huicochea, F., Velmahos, G., Cornwell, E. E., Belzberg, H., Murray, J., & Asensio, J. A. (1998). Relative bradycardia in patients with traumatic hypotension. J Trauma, 45(3), 534–539. 10.1097/00005373-199809000-00020

Dhaibar, H., Gautier, N. M., Chernyshev, O. Y., Dominic, P., & Glasscock, E. (2019). Cardiorespiratory profiling reveals primary breathing dysfunction in Kcna1-null mice: Implications for sudden unexpected death in epilepsy. Neurobiol Dis, 127, 502–511. 10.1016/j.nbd.2019.04.006

Dutsch, M., Hilz, M. J., & Devinsky, O. (2006). Impaired baroreflex function in temporal lobe epilepsy. J Neurol, 253(10), 1300–1308. 10.1007/s00415-006-0210-3

Eggleston, K. S., Olin, B. D., & Fisher, R. S. (2014). Ictal tachycardia: the head-heart connection. Seizure, 23(7), 496–505. 10.1016/j.seizure.2014.02.012

Fazan, R., Jr., de Oliveira, M., Oliveira, J. A., Salgado, H. C., & Garcia-Cairasco, N. (2011). Changes in autonomic control of the cardiovascular system in the Wistar audiogenic rat (WAR) strain. Epilepsy Behav, 22(4), 666–670. 10.1016/j.yebeh.2011.09.010

Fisher, L. A., Jessen, G., & Brown, M. R. (1983). Corticotropin-releasing factor (CRF): mechanism to elevate mean arterial pressure and heart rate. Regul Pept, 5(2), 153–161. 10.1016/0167-0115(83)90123-4

Fozard, J. R. (1983). Failure of 5-methoxytryptamine to evoke the Bezold-Jarisch effect supports homology of excitatory 5-HT receptors on vagal afferents and postganglionic sympathetic neurones. Eur J Pharmacol, 95(3-4), 331–332. 10.1016/0014-2999(83)90659-3

Gao, Y., Zhou, J. J., Zhu, Y., Kosten, T., & Li, D. P. (2017). Chronic Unpredictable Mild Stress Induces Loss of GABA Inhibition in Corticotrophin-Releasing Hormone-Expressing Neurons through NKCC1 Upregulation. Neuroendocrinology, 104(2), 194–208. 10.1159/000446114

Griffioen, K. J., Gorini, C., Jameson, H., & Mendelowitz, D. (2007). Purinergic P2X receptors mediate excitatory transmission to cardiac vagal neurons in the nucleus ambiguus after hypoxia. Hypertension, 50(1), 75–81. 10.1161/HYPERTENSIONAHA.106.086140

Gu, J., Shao, W., Liu, L., Wang, Y., Yang, Y., Zhang, Z., Wu, Y., Xu, Q., Gu, L., Zhang, Y., Shen, Y., Zhao, H., Zeng, C., & Zhang, H. (2024). Challenges and future directions of SUDEP models. Lab Anim (NY), 53(9), 226–243. 10.1038/s41684-024-01426-y

Hampel, K. G., Thijs, R. D., Elger, C. E., & Surges, R. (2017). Recurrence risk of ictal asystole in epilepsy. Neurology, 89(8), 785–791. 10.1212/WNL.0000000000004266

Heesch, C. M. (1999). Reflexes that control cardiovascular function. Am J Physiol, 277(6 Pt 2), S234–243. 10.1152/advances.1999.277.6.S234

Heusser, K., Heusser, R., Jordan, J., Urechie, V., Diedrich, A., & Tank, J. (2021). Baroreflex Curve Fitting Using a WYSIWYG Boltzmann Sigmoidal Equation. Front Neurosci, 15, 697582. 10.3389/fnins.2021.697582

Hooper, A., Fuller, P. M., & Maguire, J. (2018). Hippocampal corticotropin-releasing hormone neurons support recognition memory and modulate hippocampal excitability. PLoS One, 13(1), e0191363. 10.1371/journal.pone.0191363

Hsueh, B., Chen, R., Jo, Y., Tang, D., Raffiee, M., Kim, Y. S., Inoue, M., Randles, S., Ramakrishnan, C., Patel, S., Kim, D. K., Liu, T. X., Kim, S. H., Tan, L., Mortazavi, L., Cordero, A., Shi, J., Zhao, M., Ho, T. T.,…Deisseroth, K. (2023). Cardiogenic control of affective behavioural state. Nature, 615(7951), 292–299. 10.1038/s41586-023-05748-8

Jackson, I. P., Heppler, M., & Highley, A. D. (2024). BEZOLD-JARISCH REFLEX IN THE SETTING OF NEUROGENIC AND CARDIOGENIC SHOCK. CHEST, 166(4), A2491–A2492. 10.1016/j.chest.2024.06.1517

Jarisch, A., & Richter, H. (1939). Die afferenten Bahnen des Veratrineffektes in den Herznerven. Naunyn-Schmiedebergs Archiv für experimentelle Pathologie und Pharmakologie, 193(2), 355–371.

Jeppesen, J., Fuglsang-Frederiksen, A., Johansen, P., Christensen, J., Wustenhagen, S., Tankisi, H., Qerama, E., Hess, A., & Beniczky, S. (2019). Seizure detection based on heart rate variability using a wearable electrocardiography device. Epilepsia, 60(10), 2105–2113. 10.1111/epi.16343

Karlen-Amarante, M., da Cunha, N. V., de Andrade, O., de Souza, H. C., & Martins-Pinge, M. C. (2012). Altered baroreflex and autonomic modulation in monosodium glutamate-induced hyperadipose rats. Metabolism, 61(10), 1435–1442. 10.1016/j.metabol.2012.03.005

Kaya, C. A., Ozkaynakci, A. E., Goren, M. Z., & Onat, F. Y. (2005). Changes in baroreflex responses of kindled rats. Epilepsia, 46(3), 367–371. 10.1111/j.0013-9580.2005.32904.x

Kinney, H. C., Richerson, G. B., Dymecki, S. M., Darnall, R. A., & Nattie, E. E. (2009). The brainstem and serotonin in the sudden infant death syndrome. Annu Rev Pathol, 4, 517–550. 10.1146/annurev.pathol.4.110807.092322

Kuipers, J. R., Sidi, D., Heymann, M. A., & Rudolph, A. M. (1984). Effects of nitroprusside on cardiac function, blood flow distribution, and oxygen consumption in the conscious young lamb. Pediatr Res, 18(7), 618–626. 10.1203/00006450-198407000-00010

Lamrani, Y., Tran, T. P. Y., Toffa, D. H., Robert, M., Berube, A. A., Nguyen, D. K., & Bou Assi, E. (2023). Unexpected cardiorespiratory findings postictally and at rest weeks prior to SUDEP. Front Neurol, 14, 1129395. 10.3389/fneur.2023.1129395

Lee, S. K., Ryu, P. D., & Lee, S. Y. (2013). Differential distributions of neuropeptides in hypothalamic paraventricular nucleus neurons projecting to the rostral ventrolateral medulla in the rat. Neurosci Lett, 556, 160–165. 10.1016/j.neulet.2013.09.070

Little, R. A., Kirkman, E., Driscoll, P., Hanson, J., & Mackway-Jones, K. (1995). Preventable deaths after injury: why are the traditional ‘vital’ signs poor indicators of blood loss? J Accid Emerg Med, 12(1), 1–14. 10.1136/emj.12.1.1

Lovelace, J. W., Ma, J., Yadav, S., Chhabria, K., Shen, H., Pang, Z., Qi, T., Sehgal, R., Zhang, Y., Bali, T., Vaissiere, T., Tan, S., Liu, Y., Rumbaugh, G., Ye, L., Kleinfeld, D., Stringer, C., & Augustine, V. (2023). Vagal sensory neurons mediate the Bezold-Jarisch reflex and induce syncope. Nature, 623(7986), 387–396. 10.1038/s41586-023-06680-7

Lovick, T. A., & Coote, J. H. (1988). Electrophysiological properties of paraventriculo-spinal neurones in the rat. Brain Res, 454(1-2), 123–130. 10.1016/0006-8993(88)90810-4

Mackay, M., Mahlaba, H., Gavillet, E., & Whittaker, R. G. (2017). Seizure self-prediction: Myth or missed opportunity? Seizure, 51, 180–185. 10.1016/j.seizure.2017.08.011

Manka-Gaca, I., Labuz-Roszak, B., Machowska-Majchrzak, A., Kalarus, Z., Sredniawa, B., & Pierzchala, K. (2016). Interictal heart rate in patients with epilepsy. Wiad Lek, *69*(3 pt 2), 443–448.

Mayer, H., Benninger, F., Urak, L., Plattner, B., Geldner, J., & Feucht, M. (2004). EKG abnormalities in children and adolescents with symptomatic temporal lobe epilepsy. Neurology, 63(2), 324–328. 10.1212/01.wnl.0000129830.72973.56

Melon, L. C., Hooper, A., Yang, X., Moss, S. J., & Maguire, J. (2018). Inability to suppress the stress-induced activation of the HPA axis during the peripartum period engenders deficits in postpartum behaviors in mice. Psychoneuroendocrinology, 90, 182–193. 10.1016/j.psyneuen.2017.12.003

Moga, M. M., Saper, C. B., & Gray, T. S. (1990). Neuropeptide organization of the hypothalamic projection to the parabrachial nucleus in the rat. J Comp Neurol, 295(4), 662–682. 10.1002/cne.902950409

Moore, B. M., Jerry Jou, C., Tatalovic, M., Kaufman, E. S., Kline, D. D., & Kunze, D. L. (2014). The Kv1.1 null mouse, a model of sudden unexpected death in epilepsy (SUDEP). Epilepsia, 55(11), 1808–1816. 10.1111/epi.12793

Moseley, B. D., Ghearing, G. R., Benarroch, E. E., & Britton, J. W. (2011). Early seizure termination in ictal asystole. Epilepsy Res, 97(1-2), 220–224. 10.1016/j.eplepsyres.2011.08.008

Mueller, S. G., Nei, M., Bateman, L. M., Knowlton, R., Laxer, K. D., Friedman, D., Devinsky, O., & Goldman, A. M. (2018). Brainstem network disruption: A pathway to sudden unexplained death in epilepsy? Hum Brain Mapp, 39(12), 4820–4830. 10.1002/hbm.24325

Murugesan, A., Rani, M. R. S., Hampson, J., Zonjy, B., Lacuey, N., Faingold, C. L., Friedman, D., Devinsky, O., Sainju, R. K., Schuele, S., Diehl, B., Nei, M., Harper, R. M., Bateman, L. M., Richerson, G., & Lhatoo, S. D. (2018). Serum serotonin levels in patients with epileptic seizures. Epilepsia, 59(6), e91–e97. 10.1111/epi.14198

Nakase, K., Kollmar, R., Lazar, J., Arjomandi, H., Sundaram, K., Silverman, J., Orman, R., Weedon, J., Stefanov, D., Savoca, E., Tordjman, L., Stiles, K., Ihsan, M., Nunez, A., Guzman, L., & Stewart, M. (2016). Laryngospasm, central and obstructive apnea during seizures: Defining pathophysiology for sudden death in a rat model. Epilepsy Res, 128, 126–139. 10.1016/j.eplepsyres.2016.08.004

Nass, R. D., Hampel, K. G., Elger, C. E., & Surges, R. (2019). Blood Pressure in Seizures and Epilepsy. Front Neurol, 10, 501. 10.3389/fneur.2019.00501

Nijsen, M. J., Croiset, G., Stam, R., Bruijnzeel, A., Diamant, M., de Wied, D., & Wiegant, V. M. (2000). The role of the CRH type 1 receptor in autonomic responses to corticotropin- releasing hormone in the rat. Neuropsychopharmacology, 22(4), 388–399. 10.1016/S0893-133X(99)00126-8

O’Toole, K. K., Hooper, A., Wakefield, S., & Maguire, J. (2014). Seizure-induced disinhibition of the HPA axis increases seizure susceptibility. Epilepsy Res, 108(1), 29–43. 10.1016/j.eplepsyres.2013.10.013

Oberg, B., & Thoren, P. (1972). Increased activity in left ventricular receptors during hemorrhage or occlusion of caval veins in the cat. A possible cause of the vaso-vagal reaction. Acta Physiol Scand, 85(2), 164–173. 10.1111/j.1748-1716.1972.tb05247.x

Ou, C. H., Tsou, M. Y., Ting, C. K., Chiou, C. S., Chan, K. H., & Tsai, S. K. (2004). Occurrence of the Bezold-Jarisch reflex during Cesarean section under spinal anesthesia—a case report. Acta Anaesthesiol Taiwan, 42(3), 175–178.

Palkovits, M. (2000). Stress-induced expression of co-localized neuropeptides in hypothalamic and amygdaloid neurons. Eur J Pharmacol, 405(1-3), 161–166. 10.1016/s0014-2999(00)00549-5

Pathak, S. J., Yousaf, M. I. K., & Shah, V. B. (2025). Sudden Unexpected Death in Epilepsy. In StatPearls.

Paulhus, K., & Glasscock, E. (2025). Seizures and premature death in mice with targeted Kv1.1 deficiency in corticolimbic circuits. Brain Commun, 7(1), fcae444. 10.1093/braincomms/fcae444

Paxinos, G., & Franklin, K. B. J. (2001). The mouse brain in stereotaxic coordinates (2nd ed.). Academic Press.

Persson, H., Ericson, M., & Tomson, T. (2003). Carbamazepine affects autonomic cardiac control in patients with newly diagnosed epilepsy. Epilepsy Res, 57(1), 69–75. 10.1016/j.eplepsyres.2003.10.012

Poulsen, D. M., & Jellinge, M. E. (2024). The Bezold-Jarisch reflex and anaesthesia. Ugeskr Laeger, 186(16). 10.61409/V11230702 (Original work published The Bezold-Jarisch reflex and anaesthesia.)

Richerson, G. B., & Buchanan, G. F. (2011). The serotonin axis: Shared mechanisms in seizures, depression, and SUDEP. Epilepsia, 52 Suppl 1(Suppl 1), 28–38. 10.1111/j.1528-1167.2010.02908.x

Rossetti, A. O., Dworetzky, B. A., Madsen, J. R., Golub, O., Beckman, J. A., & Bromfield, E. B. (2005). Ictal asystole with convulsive syncope mimicking secondary generalisation: a depth electrode study. J Neurol Neurosurg Psychiatry, 76(6), 885–887. 10.1136/jnnp.2004.051839

Rugg-Gunn, F. J., Simister, R. J., Squirrell, M., Holdright, D. R., & Duncan, J. S. (2004). Cardiac arrhythmias in focal epilepsy: a prospective long-term study. Lancet, 364(9452), 2212–2219. 10.1016/S0140-6736(04)17594-6

Ruyle, B. C., Lima-Silveira, L., Martinez, D., Cummings, K. J., Heesch, C. M., Kline, D. D., & Hasser, E. M. (2023). Paraventricular nucleus projections to the nucleus tractus solitarii are essential for full expression of hypoxia-induced peripheral chemoreflex responses. J Physiol, 601(19), 4309–4336. 10.1113/JP284907

Ryvlin, P., Nashef, L., Lhatoo, S. D., Bateman, L. M., Bird, J., Bleasel, A., Boon, P., Crespel, A., Dworetzky, B. A., Hogenhaven, H., Lerche, H., Maillard, L., Malter, M. P., Marchal, C., Murthy, J. M., Nitsche, M., Pataraia, E., Rabben, T., Rheims, S.,…Tomson, T. (2013). Incidence and mechanisms of cardiorespiratory arrests in epilepsy monitoring units (MORTEMUS): a retrospective study. Lancet Neurol, 12(10), 966–977. 10.1016/S1474-4422(13)70214-X

Ryvlin, P., Nashef, L., & Tomson, T. (2013). Prevention of sudden unexpected death in epilepsy: a realistic goal? Epilepsia, 54 *Suppl 2*, 23–28. 10.1111/epi.12180

Salah, H. M., Gupta, R., Hicks, A. J., 3rd, Mahmood, K., Haglund, N. A., Bindra, A. S., Antoine, S. M., Garcia, R., Yehya, A., Yaranov, D. M., Patel, P. P., Feliberti, J. P., Rollins, A. T., Rao, V. N., Letarte, L., Raje, V., Alam, A. H., Mc, C. P., Raval, N. Y.,…Fudim, M. (2025). Baroreflex Function in Cardiovascular Disease. J Card Fail, 31(1), 117–126. 10.1016/j.cardfail.2024.08.062

San Antonio-Arce, V., Konig, A. K., Klotz, K. A., Schonberger, J., Schulze-Bonhage, A., & Jacobs-Le Van, J. (2024). Ictal tachycardia in children with epilepsy. Seizure, 123, 128–132. 10.1016/j.seizure.2024.11.007

Savic, B., Murphy, D., & Japundzic-Zigon, N. (2022). The Paraventricular Nucleus of the Hypothalamus in Control of Blood Pressure and Blood Pressure Variability. Front Physiol, 13, 858941. 10.3389/fphys.2022.858941

Sawyer, N. T., & Escayg, A. (2010). Stress and epilepsy: multiple models, multiple outcomes. J Clin Neurophysiol, 27(6), 445–452. 10.1097/WNP.0b013e3181fe0573

Schadt, J. C. (1989). Sympathetic and hemodynamic adjustments to hemorrhage: a possible role for endogenous opioid peptides. Resuscitation, 18(2-3), 219–228. 10.1016/0300-9572(89)90024-5

Schreihofer, A. M., & Guyenet, P. G. (2002). The baroreflex and beyond: control of sympathetic vasomotor tone by GABAergic neurons in the ventrolateral medulla. Clin Exp Pharmacol Physiol, 29(5-6), 514–521. 10.1046/j.1440-1681.2002.03665.x

Schuele, S. U., Afshari, M., Afshari, Z. S., Macken, M. P., Asconape, J., Wolfe, L., & Gerard, E. E. (2011). Ictal central apnea as a predictor for sudden unexpected death in epilepsy. Epilepsy Behav, 22(2), 401–403. 10.1016/j.yebeh.2011.06.036

Schuele, S. U., Bermeo, A. C., Alexopoulos, A. V., & Burgess, R. C. (2010). Anoxia-ischemia: a mechanism of seizure termination in ictal asystole. Epilepsia, 51(1), 170–173. 10.1111/j.1528-1167.2009.02168.x

Sevcencu, C., & Struijk, J. J. (2010). Autonomic alterations and cardiac changes in epilepsy. Epilepsia, 51(5), 725–737. 10.1111/j.1528-1167.2009.02479.x

Si, M., Darvish, A., Paulhus, K., Kumar, P., Hamilton, K. A., & Glasscock, E. (2024). Epilepsy-associated Kv1.1 channel subunits regulate intrinsic cardiac pacemaking in mice. J Gen Physiol, 156(9). 10.1085/jgp.202413578

So, J., Shin, W. J., & Shim, J. H. (2013). A cardiovascular collapse occurred in the beach chair position for shoulder arthroscopy under general anesthesia -A case report. Korean J Anesthesiol, 64(3), 265–267. 10.4097/kjae.2013.64.3.265

Souza, G., Stornetta, R. L., Stornetta, D. S., Guyenet, P. G., & Abbott, S. B. G. (2022). Adrenergic C1 neurons monitor arterial blood pressure and determine the sympathetic response to hemorrhage. Cell Rep, 38(10), 110480. 10.1016/j.celrep.2022.110480

Stewart, M., Kollmar, R., Nakase, K., Silverman, J., Sundaram, K., Orman, R., & Lazar, J. (2017). Obstructive apnea due to laryngospasm links ictal to postictal events in SUDEP cases and offers practical biomarkers for review of past cases and prevention of new ones. Epilepsia, 58(6), e87–e90. 10.1111/epi.13765

Strain, M. M., Espinoza, L., Fedorchak, S., Littlejohn, E. L., Andrade, M. A., Toney, G. M., & Boychuk, C. R. (2023). Early central cardiovagal dysfunction after high fat diet in a murine model. Sci Rep, 13(1), 6550. 10.1038/s41598-023-32492-w

Sutula, T. P., & Dudek, F. E. (2007). Unmasking recurrent excitation generated by mossy fiber sprouting in the epileptic dentate gyrus: an emergent property of a complex system. Prog Brain Res, 163, 541–563. 10.1016/S0079-6123(07)63029-5

Tomson, T., Walczak, T., Sillanpaa, M., & Sander, J. W. (2005). Sudden unexpected death in epilepsy: a review of incidence and risk factors. Epilepsia, 46 *Suppl 11*, 54–61. 10.1111/j.1528-1167.2005.00411.x

Trosclair, K., Dhaibar, H. A., Gautier, N. M., Mishra, V., & Glasscock, E. (2020). Neuron-specific Kv1.1 deficiency is sufficient to cause epilepsy, premature death, and cardiorespiratory dysregulation. Neurobiol Dis, 137, 104759. 10.1016/j.nbd.2020.104759

Tyagi, T., Ahmad, S., Gupta, N., Sahu, A., Ahmad, Y., Nair, V., Chatterjee, T., Bajaj, N., Sengupta, S., Ganju, L., Singh, S. B., & Ashraf, M. Z. (2014). Altered expression of platelet proteins and calpain activity mediate hypoxia-induced prothrombotic phenotype. Blood, 123(8), 1250–1260. 10.1182/blood-2013-05-501924

Verberne, A. J., & Guyenet, P. G. (1992). Medullary pathway of the Bezold-Jarisch reflex in the rat. Am J Physiol, *263*(6 Pt 2), R1195–1202. 10.1152/ajpregu.1992.263.6.R1195

Wang, L. A., Nguyen, D. H., & Mifflin, S. W. (2018). CRHR2 (Corticotropin-Releasing Hormone Receptor 2) in the Nucleus of the Solitary Tract Contributes to Intermittent Hypoxia-Induced Hypertension. Hypertension, 72(4), 994–1001. 10.1161/HYPERTENSIONAHA.118.11497

Wang, L. A., Nguyen, D. H., & Mifflin, S. W. (2019). Corticotropin-releasing hormone projections from the paraventricular nucleus of the hypothalamus to the nucleus of the solitary tract increase blood pressure. J Neurophysiol, 121(2), 602–608. 10.1152/jn.00623.2018

Whalen, E. J., Johnson, A. K., Lewis, S. J., & New Collective, A. (2000). Functional evidence for the rapid desensitization of 5-HT(3) receptors on vagal afferents mediating the Bezold-Jarisch reflex. Brain Res, 873(2), 302–305. 10.1016/s0006-8993(00)02526-9

Yamaguchi, J., Andrade, M. A., Truong, T. T., & Toney, G. M. (2024). Glutamate Spillover Dynamically Strengthens Gabaergic Synaptic Inhibition of the Hypothalamic Paraventricular Nucleus. J Neurosci, 44(7). 10.1523/JNEUROSCI.1851-22.2023

Yamano, M., Ito, H., Kamato, T., & Miyata, K. (1995). Characteristics of inhibitory effects of serotonin (5-HT)3-receptor antagonists, YM060 and YM114 (KAE-393), on the von Bezold-Jarisch reflex induced by 2-Methyl-5-HT, veratridine and electrical stimulation of vagus nerves in anesthetized rats. Jpn J Pharmacol, 69(4), 351–356. 10.1254/jjp.69.351

Zeidler, Z., Brandt-Fontaine, M., Leintz, C., Krook-Magnuson, C., Netoff, T., & Krook-Magnuson, E. (2018). Targeting the Mouse Ventral Hippocampus in the Intrahippocampal Kainic Acid Model of Temporal Lobe Epilepsy. eNeuro, 5(4). 10.1523/ENEURO.0158-18.2018

Zijlmans, M., Flanagan, D., & Gotman, J. (2002). Heart rate changes and ECG abnormalities during epileptic seizures: prevalence and definition of an objective clinical sign. Epilepsia, 43(8), 847–854. 10.1046/j.1528-1157.2002.37801.x

