## Supplemental Tables for "Autonomic reflex plasticity associates with time-dependent SUDEP susceptibility in a murine model with hyperreactive stress circuits"

| Table 1.1: Figure 1A | |
|---|---|
| Genotype | Resting HR |
| Kcc2/Crh | 532.1 ± 55.0 BPM, n=23 |
| WT | 555.8 ± 49.5 BPM, n=13 |
| Unpaired t-test | p=0.2071 |

| Table 1.2: Figure 1C- HR normalized to Day 0 (BPM) | | | | |
|---|---|---|---|---|
| | Day 7 ΔHR | Day 14 ΔHR | Day 21 ΔHR | Day 28 ΔHR |
| Kcc2/Crh +vIHKA | 43.6 ± 44.1, n=14 | 53.8 ± 32.0, n=11 | 39.2 ± 49.8, n=10 | 40.6 ± 43.8, n=9 |
| Kcc2/Crh + Saline | -11.2 ± 30.0, n=8 | -11.2 ± 30.0, n=8 | -3.1 ± 51.1, n=8 | 8.0 ± 39.6, n=8 |
| WT + vIHKA | 26.1 ± 12.1, n=6 | 37.0 ± 35.5, n=6 | 50.8 ± 45.5, n=6 | 6.7 ± 63.8, n=5 |
| WT + Saline | 5.6 ± 28.7, n=7 | -11.6 ± 18.0, n=7 | -5.6 ± 21.7, n=7 | -6.3 ± 36.4, n=6 |

| Mixed-effects Analysis | | | |
|---|---|---|---|
| Mixed-effects Analysis | **Time Factor** | **F (3.362, 92.45) = 2.833** | **p=0.0369** |
| | **Group Factor** | **F (3, 31) = 6.831** | **p=0.0012** |
| | **Interaction** | **F (10.09, 92.45) = 1.974** | **p=0.0445** |

Multiple Comparisons with Tukey's Post Hoc Test

| Comparison | Days post vIHKA or Saline 7 | 14 | 21 | 28 |
|---|---|---|---|---|
| Kcc2/Crh+vIHKA vs. Kcc2/Crh+Saline | **p=0.0127** | p=0.0523 | p=0.3280 | p=0.4009 |
| Kcc2/Crh +vIHKA vs. WT+vIHKA | p=0.5370 | p=0.7696 | p=0.9631 | p=0.7245 |
| Kcc2/Crh+vIHKA vs. WT+Saline | p=0.1202 | **p=0.0002** | p=0.1021 | p=0.1641 |
| Kcc2/Crh+Saline vs. WT+vIHKA | **p=0.0415** | p=0.5540 | p=0.2171 | p>0.9999 |
| Kcc2/Crh+Saline vs. WT+Saline | p=0.6905 | p=0.2903 | p=0.9992 | p=0.8958 |
| WT+vIHKA vs. WT+Saline | p=0.3712 | p=0.0702 | p=0.1013 | p=0.9759 |

| Comparison | Kcc2/Crh + vIHKA | Kcc2/Crh + Saline | WT+ vIHKA | WT+ Saline |
|---|---|---|---|---|
| Day 0 vs. Day 7 | **p=0.0190** | p=0.8201 | **p=0.0169** | p=0.9828 |
| Day 0 vs. Day 14 | **p=0.0017** | p=0.7795 | p=0.2159 | p=0.4952 |
| Day 0 vs. Day 21 | p=0.1763 | p=0.9998 | p=0.1784 | p=0.9534 |
| Day 0 vs. Day 28 | p=0.1253 | p=0.9757 | p=0.999 | p=0.9912 |
| Day 7 vs. Day 14 | p=0.4912 | p=0.5314 | p=0.9521 | p=0.7076 |
| Day 7 vs. Day 21 | p>0.9999 | p=0.9962 | p=0.6919 | p=0.8957 |
| Day 7 vs. Day 28 | p=0.9983 | p=0.8873 | p=0.8843 | p=0.9938 |
| Day 14 vs. Day 21 | p=0.7220 | p=0.7816 | p=0.7776 | p=0.9703 |
| Day 14 vs. Day 28 | p=0.3947 | p=0.9991 | p=0.9372 | p=0.9997 |
| Day 21 vs. Day 28 | p=0.9957 | p=0.9885 | p=0.6683 | p>0.9999 |

| Table 2.1: Figure 2 | | |
|---|---|---|
| Parameter | Genotype | Value |
| Fig 2D. Pre-ictal HR change onset (sec before seizure onset) | Kcc2/Crh + vIHKA | -7.0 ± 7.2 sec, n=7 |
| | WT + vIHKA | -8.2 ± 5.4 sec, n=5 |
| | unpaired t-test | p=0.7607 |
| Fig 2E. Average ictal HR, relative to baseline | Kcc2/Crh + vIHKA | 3.91 ± 87.92 BPM, n=8 |
| | WT + vIHKA | 36.75 ± 59.80 BPM, n=6 |
| | unpaired t-test | p=0.4476 |
| Fig 2F. Average post-ictal HR, relative to baseline (0-10 sec) | Kcc2/Crh + vIHKA | 2.0 ± 88.4 BPM, n=8 |
| | WT + vIHKA | 10.2± 58.3 BPM, n=6 |
| | unpaired t-test | p=0.8453 |
| Fig 2F. Average post-ictal HR, relative to baseline (10-30 sec) | Kcc2/Crh + vIHKA | 43.7 ± 84.6 BPM, n=8 |
| | WT + vIHKA | 79.8 ± 65.1 BPM, n=6 |
| | unpaired t-test | p=0.4029 |
| Fig 2E. Magnitude Ictal Tachycardia, ΔHR | Kcc2/Crh + vIHKA | 65.6 ± 54.9 BPM, n=8 |
| | WT + vIHKA | 75.2 ± 61.72 BPM, n=6 |
| | unpaired t-test | p=0.7629 |
| Fig 2F. Duration Ictal Tachycardia | Kcc2/Crh + vIHKA | 16.3 ± 20.0 sec, n=8 |
| | WT + vIHKA | 13.5 ± 12.3 sec, n=6 |
| | unpaired t-test | p=0.7721 |
| **Fig 2I. Magnitude Ictal Bradycardia, ΔHR** | Kcc2/Crh + vIHKA | -208.1 ± 63.5 BPM, n=6 |
| | WT + vIHKA | -113.2 ± 53.0 BPM, n=5 |
| | **unpaired t-test** | **p=0.0264** |
| **Fig 2J. Duration Ictal Bradycardia** | Kcc2/Crh + vIHKA | 11.6 ±3.3 sec, n=6 |
| | WT + vIHKA | 7.2 ± 2.2 sec, n=5 |
| | **unpaired t-test** | **p=0.0350** |
| Seizure Duration (not shown) | Kcc2/Crh + vIHKA | 31.3 ±13.4 sec, n=8 |
| | WT + vIHKA | 38.5 ±9.5 sec, n=6 |
| | unpaired t-test | p=0.2871 |
| Magnitude Pre-ictal Tachycardia, ΔHR (not shown) | Kcc2/Crh + vIHKA | 77.8 ± 56.2 BPM, n=6 |
| | WT + vIHKA | 62.4 ± 24.5 BPM, n=4 |
| | unpaired t-test | p=0.6245 |

| Table 3.1: Figure 3C- Phenylephrine mediated increase in BP (ΔmmHg) | | | | |
|---|---|---|---|---|
| | | Time | | |
| | | Saline Injected | 10 days post vIHKA | 30 days post vIHKA |
| Genotype | Kcc2/Crh | 91.7 ± 14.7, n=5 | 89.3 ± 17.6, n=5 | 87.1 ± 12.4, n=6 |
| | WT | 81.1 ± 17.4, n=5 | 95.2 ± 8.4, n=5 | 104.2 ± 11.6, n=5 |
| Two-way ANOVA | | Interaction | F (2, 25) = 2.571 | p=0.0965 |
| | | Time Factor | F (2, 25) = 1.150 | p=0.3327 |
| | | Genotype Factor | F (1, 25) = 0.6718 | p=0.4202 |

| Table 3.2: Figure 3E- Slope (ΔBPM/ΔmmHg) | | | | |
|---|---|---|---|---|
| | | Time | | |
| | | Saline Injected | 10 days post vIHKA | 30 days post vIHKA |
| Genotype | Kcc2/Crh | -14.7 ± 7.6, n=5 | -18.5 ± 5.0, n=5 | -11.5 ± 9.8, n=6 |
| | WT | -9.2 ± 8.5, n=5 | -16.0 ± 7.1, n=5 | -4.2 ± 4.0, n=5 |
| Two-way ANOVA | | Interaction | F (2, 25) = 0.2824 | p=0.7564 |
| | | **Time Factor** | **F (2, 25) = 4.243** | **p=0.0259** |
| | | Genotype Factor | F (1, 25) = 3.731 | p=0.0648 |
| Tukey's Multiple Comparisons | | Time (Main Row Effect) | saline vs. 10 | p=0.2593 |
| | | | saline vs. 30 | p=0.4287 |
| | | | **10 vs. 30** | **p=0.0198** |

| Table 3.3: Figure 3D- V50 (mmHg) | | | | |
|---|---|---|---|---|
| | | Time | | |
| | | Saline Injected | 10 days post vIHKA | 30 days post vIHKA |
| Genotype | Kcc2/Crh | 115.5 ± 14.1, n=5 | 114.9 ± 12.9, n=5 | 126.3± 14.4, n=6 |
| | WT | 118.2 ± 15.8, n=5 | 116.4 ± 10.8, n=5 | 138.2 ± 17.3, n=5 |
| Two-way ANOVA | | Interaction | F (2, 25) = 0.4157 | p=0.6644 |
| | | **Time Factor** | **F (2, 25) = 4.407** | **p=0.0229** |
| | | Genotype Factor | F (1, 25) = 1.091 | p=0.3062 |
| Tukey's Multiple Comparisons | | Time (Main Row Effect) | saline vs. 10 | p=0.9803 |
| | | | saline vs. 30 | p=0.0545 |
| | | | **10 vs. 30** | **p=0.0359** |

| Table 4.1: Figure 4C | | | | |
|---|---|---|---|---|
| | | | SNP-ΔBP from baseline (mmHg) | |
| | | | Early | Late |
| Group | Kcc2/Crh | Saline | -36.3 ± 9.1, n=5 | -40.0 ± 6.8, n=5 |
| | | 10 days post vIHKA | -33.2 ± 10.9, n=5 | -35.3 ± 11.2, n=5 |
| | | 30 days post vIHKA | -37.1 ± 9.9, n=6 | -39.6 ± 9.1, n=6 |
| | WT | Saline | -40.6 ± 9.8, n=6 | -44.6 ± 9.4, n=6 |
| | | 10 days post vIHKA | -40.5 ± 10.2, n=5 | -48.2 ± 8.3, n=5 |
| | | 30 days post vIHKA | -36.1 ± 8.0, n=6 | -38.4 ± 8.1, n=6 |
| Repeated Measures Two-way ANOVA | | Interaction | F (10, 54) = 0.9737 | P=0.4767 |
| | | **Time Factor** | **F (2, 54) = 554.4** | **P<0.0001** |
| | | Group Factor | F (5, 27) = 0.8506 | P=0.5263 |
| Dunnett's Multiple Comparisons | | Time (Main Row Effect) | baseline vs. early | **p<0.0001** |
| | | | baseline vs. late | **p<0.0001** |

| Table 4.2: Figure 4D | | | | |
|---|---|---|---|---|
| | | | SNP-ΔHR from baseline (BPM) | |
| | | | Early | Late |
| Group | Kcc2/Crh | Saline | 5.3 ± 7.5, n=5 | 10.6 ± 10.3, n=5 |
| | | 10 days post vIHKA | 4.4 ± 5.7, n=5 | -15.9 ± 21.9, n=5 |
| | | 30 days post vIHKA | -7.5 ± 11.9, n=6 | -32.9 ± 31.7, n=6 |
| | WT | Saline | 0.7 ± 2.7, n=6 | -6.3 ± 8.5, n=6 |
| | | 10 days post vIHKA | -1.1 ± 6.9, n=5 | -31.3 ± 27.7, n=5 |
| | | 30 days post vIHKA | 3.0 ± 16.2, n=6 | -2.4 ± 18.6, n=6 |
| Repeated Measures Two-way ANOVA | | **Interaction** | **F (10, 54) = 3.327** | **P=0.0020** |
| | | **Time Factor** | **F (2, 54) = 14.78** | **P<0.0001** |
| | | **Group Factor** | **F (5, 27) = 2.721** | **P=0.0408** |
| Multiple comparisons with Tukey's Post Hoc | | | | |
| | | Saline | 10 | 30 |
| Kcc2/Crh | baseline vs. early | p=0.7524 | p=0.8220 | p=0.4998 |
| | baseline vs. late | p=0.3240 | p=0.0853 | **p<0.0001** |
| | early vs. late | p=0.7485 | **p=0.0206** | **p=0.0010** |
| WT | baseline vs. early | p=0.9944 | p=0.9868 | p=0.8921 |
| | baseline vs. late | p=0.6153 | **p=0.0002** | p=0.9297 |
| | early vs. late | p=0.5523 | **p=0.0004** | p=0.6926 |
| Comparison | | | Early | Late |
| WT + Saline vs. Kcc2/Crh + Saline | | | p=0.9936 | p=0.3309 |
| WT + Saline vs. WT + vIHKA day 10 | | | p>0.9999 | **p=0.0384** |
| WT + Saline vs. WT + vIHKA day 30 | | | p=0.9997 | p=0.9964 |
| WT + Saline vs. Kcc2/Crh + vIHKA day 10 | | | p=0.9977 | p=0.8561 |
| WT + Saline vs. Kcc2/Crh + vIHKA day 30 | | | p=0.9030 | **p=0.0141** |
| WT + vIHKA day 10 vs. WT +vIHKA day 30 | | | p=0.9959 | **p=0.0100** |
| WT + vIHKA day 10 vs. Kcc2/Crh + Saline | | | p=0.9763 | **p<0.0001** |
| WT + vIHKA day 10 vs. Kcc2/Crh + vIHKA day 10 | | | p=0.988 | p=0.4821 |
| WT + vIHKA day 10 vs. Kcc2/Crh + vIHKA day 30 | | | p=0.9712 | p>0.9999 |
| Kcc2/Crh + Saline vs. WT + vIHKA day 30 | | | p=0.9998 | p=0.6202 |
| Kcc2/Crh + Saline vs. Kcc2/Crh + vIHKA day 10 | | | p>0.9999 | **p=0.0346** |
| Kcc2/Crh + Saline vs. Kcc2/Crh + vIHKA day 30 | | | p=0.6359 | **p<0.0001** |
| Kcc2/Crh + vIHKA day 10 vs. WT + vIHKA day 30 | | | p>0.9999 | p=0.586 |
| Kcc2/Crh + vIHKA day 10 vs. Kcc2/Crh + vIHKA day 30 | | | p=0.7043 | p=0.3189 |
| Kcc2/Crh + vIHKA day 30 vs. WT + vIHKA day 30 | | | p=0.7622 | **p=0.0030** |

| Table 4.3: Figure 4E | | | | |
|---|---|---|---|---|
| | | | SNP mediated ΔHR/ΔBP | |
| | | | Early | Late |
| Group | Kcc2/Crh | Saline | -0.14 ± 0.22, n=5 | -0.25 ± 0.27, n=5 |
| | | 10 days post vIHKA | -0.10 ± 0.20, n=5 | 0.74 ± 1.28, n=5 |
| | | 30 days post vIHKA | 0.17 ± 0.34, n=6 | 0.76 ± 0.65, n=6 |
| | WT | Saline | -0.01 ± 0.07, n=6 | 0.15 ± 0.19, n=6 |
| | | 10 days post vIHKA | 0.03± 0.18, n=5 | 0.65 ± 0.57, n=5 |
| | | 30 days post vIHKA | -0.10 ± 0.52, n=6 | 0.05 ± 0.51, n=6 |
| Repeated Measures Two-Way ANOVA | | **Interaction** | **F (5, 27) = 2.612** | **p=0.0473** |
| | | **Time Factor** | **F (1, 27) = 17.60** | **p=0.0003** |
| | | Group Factor | F (5, 27) = 1.737 | p=0.1602 |
| Multiple comparisons with Tukey's Post hoc | | | | |
| Group | Kcc2/Crh | Saline | early vs. late | p=0.6380 |
| | | **10 days post vIHKA** | **early vs. late** | **p=0.0011** |
| | | **30 days post vIHKA** | **early vs. late** | **p=0.0094** |
| | WT | Saline | early vs. late | p=0.4484 |
| | | **10 days post vIHKA** | **early vs. late** | **p=0.0106** |
| | | 30 days post vIHKA | early vs. late | p=0.4868 |
| Comparison | | | Early | Late |
| WT + Saline vs. Kcc2/Crh + Saline | | | p=0.9983 | p=0.7944 |
| WT + Saline vs. WT + vIHKA day 10 | | | p>0.9999 | p=0.5982 |
| WT + Saline vs. WT + vIHKA day 30 | | | p=0.9996 | p=0.9992 |
| WT + Saline vs. Kcc2/Crh + vIHKA day 10 | | | p=0.9997 | p=0.4212 |
| WT + Saline vs. Kcc2/Crh + vIHKA day 30 | | | p=0.9894 | p=0.3335 |
| WT + vIHKA day 10 vs. WT +vIHKA day 30 | | | p=0.9985 | p=0.3845 |
| WT + vIHKA day 10 vs. Kcc2/Crh + Saline | | | p=0.9960 | p=0.0793 |
| WT + vIHKA day 10 vs. Kcc2/Crh + vIHKA day 10 | | | p=0.9989 | p=0.9998 |
| WT + vIHKA day 10 vs. Kcc2/Crh + vIHKA day 30 | | | p=0.9966 | p=0.9994 |
| Kcc2/Crh + Saline vs. WT + vIHKA day 30 | | | p>0.9999 | p=0.9350 |
| Kcc2/Crh + Saline vs. Kcc2/Crh + vIHKA day 10 | | | p>0.9999 | **p=0.0411** |
| Kcc2/Crh + Saline vs. Kcc2/Crh + vIHKA day 30 | | | p=0.9150 | **p=0.0247** |
| Kcc2/Crh + vIHKA day 10 vs. WT + vIHKA day 30 | | | p>0.9999 | p=0.2414 |
| Kcc2/Crh + vIHKA day 10 vs. Kcc2/Crh + vIHKA day 30 | | | p=0.9496 | p>0.9999 |
| Kcc2/Crh + vIHKA day 30 vs. WT + vIHKA day 30 | | | p=0.9367 | p=0.1743 |

| Table 5.1: Figure 5B- PBG mediated ΔHR (BPM) | | | |
|---|---|---|---|
| | Drug | | |
| Genotype | Baseline | + atenolol | +Scopolamine |
| WT | -251.9 ± 155.5, n=6 | -286.6 ± 90.9, n=6 | -42.0 ± 9.8, n=6 |
| RM One-way ANOVA | **Treatment** | **F (1.282, 6.409)= 15.76** | **p=0.0051** |
| Multiple Comparisons Tukey's Post Hoc | | Baseline vs. Atenolol | p=0.6729 |
| | | **Baseline vs. Scopolamine** | **p=0.0441** |
| | | **Atenolol vs. Scopolamine** | **p=0.0023** |

| Table 5.2: Figure 5E- PBG mediated ΔHR over time (BPM) | | | | |
|---|---|---|---|---|
| | | Drug | | |
| | | Saline Injected (ΔHR) | 10 days post vIHKA (ΔHR) | 30 days post vIHKA (ΔHR) |
| Genotype | Kcc2/Crh | -189.7 ± 119.9, n=11 | -390.3 ± 77.1, n=9 | -222.8 ± 114.9, n=8 |
| | WT | -181.8 ± 111.5, n=7 | -183.0 ± 123.1, n=6 | -120.4 ± 92.9; n=6 |
| Two-way ANOVA | **Interaction** | | **F (2, 41) = 3.343** | **P=0.0452** |
| | **Time Factor** | | **F (2, 41) = 4.909** | **P=0.0123** |
| | **Genotype Factor** | | **F (1, 41) = 10.78** | **P=0.0021** |
| Multiple Comparisons with Tukey's Post Hoc | Time | Saline | WT vs. Kcc2/Crh | P= 0.8814 |
| | | **10 days post vIHKA** | **WT vs. Kcc2/Crh** | **P= 0.0007** |
| | | 30 days post vIHKA | WT vs. Kcc2/Crh | P= 0.0864 |
| | Genotype | WT | saline vs. 10 | P= 0.9998 |
| | | | saline vs. 30 | P= 0.5666 |
| | | | 10 vs. 30 | P= 0.5782 |
| | | Kcc2/Crh | **saline vs. 10** | **P= 0.0005** |
| | | | saline vs. 30 | P = 0.7874 |
| | | | **10 vs. 30** | **P= 0.0074** |

| Table 6.1: Figure 6B- Dentate Granule Cell Density (# cells/area) | | | | |
|---|---|---|---|---|
| | | Time | | |
| | | Saline Injected | 10 days post vIHKA | 30 days post vIHKA |
| Genotype | Kcc2/Crh | 1.20 ± 0.11, n=4 | 0.85 ± 0.09, n=4 | 0.57 ± 0.13, n=10 |
| | WT | 1.16 ± 0.18, n=6 | 0.76 ± 0.12, n=5 | 0.56 ± 0.06, n=5 |
| Two-way ANOVA | | Interaction | F (2, 28) = 0.2357 | p=0.7916 |
| | | **Time Factor** | **F (2, 28) = 66.34** | **p<0.0001** |
| | | Genotype Factor | F (1, 28) = 1.206 | p=0.2816 |
| Tukey's Multiple Comparisons | | Time (Main Row Effect) | **saline vs. 10** | **p<0.0001** |
| | | | **saline vs. 30** | **p<0.0001** |
| | | | **10 vs. 30** | **p=0.0005** |

| Table 6.2: Figure 6C- Mossy Fiber Sprouting (Mean Fluorescence Intensity) | | | | |
|---|---|---|---|---|
| | | Time | | |
| | | Saline Injected | 10 days post vIHKA | 30 days post vIHKA |
| Genotype | Kcc2/Crh | 12.4 ± 3.1, n=4 | 22.2± 1.3, n=4 | 19.4 ± 2.9, n=10 |
| | WT | 10.6 ± 4.2, n=6 | 17.9 ± 2.2, n=5 | 21.2 ± 4.0, n=5 |
| Two-way ANOVA | | Interaction | F (2, 28) = 2.671 | P=0.0867 |
| | | **Time Factor** | **F (2, 28) = 25.47** | **P<0.0001** |
| | | Genotype Factor | F (1, 28) = 1.614 | P=0.2144 |
| Tukey's Multiple Comparisons | | Time (Main Row Effect) | **saline vs. 10** | **p<0.0001** |
| | | | **saline vs. 30** | **p<0.0001** |
| | | | 10 vs. 30 | p=0.9834 |
